# The anti-retroviral therapy emtricitabine affects skeletal muscle DNA methylation and transcriptome patterns in a male HIV mouse model

**DOI:** 10.1101/2025.11.10.686758

**Authors:** Alok Tripathi, Shabiha Sultana, Husam Bensreti, Jie Chen, Arindam Paul, Huidong Shi, Apeksha Anand, Bharati Mendhe, Kanglun Yu, William E. Long, Rodger D. MacArthur, Eric J. Belin de Chantemele, Mark W. Hamrick, Meghan E. McGee-Lawrence

**Affiliations:** Department of Cellular Biology and Anatomy, Medical College of Georgia at Augusta University, Augusta, GA; Department Biostatistics, Data Science & Epidemiology, School of Public Health, Augusta University; Georgia Cancer Center, Medical College of Georgia at Augusta University, Augusta, GA; Department of Medicine, Medical College of Georgia at Augusta University, Augusta, GA; Vascular Biology Center, Medical College of Georgia at Augusta University, Augusta, GA

**Keywords:** DNA methylation, HIV, ART, skeletal muscle, transcriptome

## Abstract

**Background:** Although antiretroviral therapies (ART) have substantially reduced HIV-associated mortality, the increased lifespan achieved by widespread ART deployment has revealed that HIV infection is linked to an unexplained earlier onset and increased incidence of aging-associated conditions like sarcopenia. Complex syndromes, like sarcopenia, often arise from a combination of genetic and environmental factors, so in this study, we investigated effects of short-term treatment with emtricitabine (2’,3’-dideoxy-5-fluoro-3’-thiacytidine; FTC), an FDA approved ART, on skeletal muscle DNA methylation patterns and transcriptome-wide responses in a male murine model of HIV phenotypic biology (Tg26 mice).

**Methods:** We treated 6 month old male Tg26 (+/−) mice or wildtype (WT) littermates on a C57BL/6 genetic background with FTC in the drinking water for one month; control groups received drinking water vehicle alone (VEH). Muscle function and body composition were measured longitudinally. Skeletal muscle methylation patterns, transcriptional changes, and histological features were quantified at sacrifice.

**Results:** Although neither gross structural nor functional muscle deficits were observed in this short-term study with ART usage, relative decreases in muscle endurance measured by hang time over the study were 2-fold more severe in the Tg26 as compared to WT mice (p=0.0453), and markers of myogenic cell maturation (*Myf5, Myf6*) were disrupted in the Tg26 HIV model as compared to WT littermates in a manner exacerbated by FTC treatment. Fat mass, measured by DXA, also tended (p=0.085) to be uniquely increased by FTC treatment in the Tg26 mice over the study. Differential methylation patterns and pathway enrichment data suggested that the presence of an HIV phenotype and exposure to an ART regimen altered the methylation status in skeletal muscle genes such as *Camk2B, Pcolce2 and Lima1* in a manner consistent with promoting eventual functional impairment in muscle. Additionally, RNAseq revealed differential gene expression profiles and key regulatory pathways including cellular differentiation, regulation of lipid metabolism, and neuroactive ligand-receptor interactions. Lipodystrophy-related genes including *LEP, ADIPOQ and PPARα* involved in fat distribution and metabolism along with skeletal genes related to regulation of muscle strength were affected by the presence of an HIV phenotype and ART treatment.

**Conclusions:** The current study provides insights into mechanisms by which a clinically relevant ART may influence DNA methylation and transcriptome changes in skeletal muscle in the context of HIV biology. The differentially regulated pathways suggest novel targets for understanding, and eventually abrogating, the harmful effects of long-term ART use in PLWH on skeletal muscle mass and function.

## 1. INTRODUCTION

Human Immunodeficiency Virus (HIV) affects millions of individuals worldwide. As of 2022, an estimated 37.7 million people globally are living with HIV, with approximately 1.5 million new infections occurring annually. The development and increased availability of effective antiretroviral therapies (ARTs) have extended the average lifespan for people living with HIV (PLWH). However, this increased lifespan has subsequently revealed that HIV infection is linked to earlier onset and increased incidence of aging-associated conditions like cardiovascular disease, neurocognitive disorders, and sarcopenia [1]. PLWH demonstrate a decrease in skeletal muscle mass and strength compared to uninfected patients [2]. ART may promote mitochondrial dysfunction and muscle damage [3], further compounding declines in muscle mass or function in PLWH. Consistent with that hypothesis, ART-naive PLWH showed decreased muscle mass after beginning an ART regimen and experienced higher odds of decreased appendicular and total muscle mass in logistic regression models [4]. Similarly, per year use of ART associated with lower appendicular lean mass in men and women with HIV in regression models [5], and initiation of ART in PLWH associated with decreased muscle density consistent with fatty infiltration [6]. However, a molecular link between sarcopenia and ART usage has not yet been conclusively identified.

Complex syndromes, like sarcopenia, often arise from a combination of genetic and environmental factors. HIV and ART are two such environmental factors known to affect tissues [7] including skeletal muscle [8]. In blood, initial stages of HIV infection accelerate epigenetic age of peripheral blood mononuclear cells, while ART partially reverses this phenomenon [9, 10]. The skeletal muscle methylation landscape is dynamic in the context of HIV, where differential methylation patterns develop between older adults with HIV and uninfected controls following exercise [8]. Previous studies showed that skeletal muscle DNA methylation is altered by age [11], and the epigenetic clock, an aging biomarker based on DNA methylation [12], suggests accelerated epigenetic aging among PLWH [9]. These changes are positioned to contribute to the early onset of frailty in PLWH on ART, but ART-induced epigenetic modifications in skeletal muscle DNA methylation patterns have not yet been reported, and data demonstrating a causative effect of ART in skeletal muscle DNA methylation dynamics is very limited.

To address this gap in knowledge, we investigated whether combination of an HIV-like phenotype and administration of an FDA approved ART, emtricitabine (also known as 2’,3’-dideoxy-5-fluoro-3’-thiacytidine, FTC), alters the skeletal muscle DNA methylation landscape and transcriptome in male mice.

## 2. METHODS

### 2.1 Study design and tissue harvest

Mice were maintained on a 12-hour light/dark cycle as approved by the Augusta University Institutional Animal Care and Use Committee. The Tg26 model, which expresses 7 of 9 HIV proteins (absence of *gag* and *pol* renders it non-infectious) was used to mimic an HIV phenotype. Tg26 mice were backcrossed to C57BL/6J, negating kidney disease that occurs in this model on an FVB background [13]. Previous studies support use of Tg26 mice for investigating HIV-associated musculoskeletal decline [14]. 6-month-old male Tg26 (+/−) mice and wildtype (WT) littermates received FTC (Cayman Chemical #16879; n=8/group) in drinking water or water vehicle (VEH; n=8/group) for 1 month. Mice were euthanized by cervical dislocation under 5% isoflurane. The quadriceps muscles were collected at sacrifice and either frozen in liquid nitrogen or fixed in 10% neutral buffered formalin for histology.

### 2.2 Body composition

Dual-energy X-ray absorptiometry scans (DXA; Kubtec PARAMETER 3D with Digimus software, KUB Technologies, Stratford, CT, USA) were collected at baseline (prior to the onset of treatment) and immediately before sacrifice. Fat and lean mass (excluding the head) were quantified with the manufacturer’s software, and percent changes in fat and lean mass over the course of study were calculated.

### 2.3 Skeletal muscle endurance

Skeletal muscle function was evaluated at baseline and immediately before sacrifice. Endurance was measured using wire hang-time testing. Each mouse underwent three trials, with a 5-minute rest period between trials to avoid fatigue. Hang time (in seconds) was normalized to body mass.

### 2.4 Quadriceps Histology

One quadriceps muscle per mouse was histologically processed, embedded in paraffin, transversely sectioned (2–3 µm thickness), and stained with hematoxylin and eosin (H&E). Stained sections were imaged at 20x magnification using a brightfield Leica DMCS microscope and a Micropublisher six camera (Qimaging). Average fiber size and Feret diameter were measured across 25 individual fibers per section via ImageJ.

### 2.5 RNA isolation and qRT-PCR analysis

RNA was extracted from a portion of one quadriceps muscle per mouse (N=3 mice/group) via TRIzol (ThermoFisher #15-596-018), as described [15]. Methodology and primer sequences are listed in **Supplementary Methods 1.1**.

### 2.6 Reduced representation bisulfite sequencing (RRBS) library preparation and bioinformatics analysis

A portion of one quadriceps muscle per mouse (5 mg tissue; N=4 mice/group) were processed for DNA methylation analysis. DNA extraction was performed via commercial kit (Quick-DNA Plus Kit Miniprep Kit, Zymo Research, Irvine, CA). Sequence reads from reduced representation bisulfite sequencing (RRBS) libraries were identified using Illumina base calling software. Analytical and methodological details are elaborated in **Supplementary Methods 1.2**.

### 2.7 Differential methylation analysis

The beta-binomial Dispersion Shrinkage for Sequencing (DSS) model was used to identify, annotate, and visualize differentially methylated regions (DMRs), defined as putative functional regions involved in transcriptional regulation of genes, and differentially methylated CpG sites (DMCs). Cytosines with read depth ≥ 5 in ≥ 2 samples were analyzed via Wald test and Benjamini-Hochberg P-value adjustment. Significantly hyper- and hypomethylated DMCs/DMRs were defined via false discovery rate (FDR) ≤ 0.05 and methylation difference ≥ 0.1 or ≤ −0.1, respectively. Reference genomes from NCBI and UCSC were used for annotation by overlapping DMCs or DMRs with functional regions (genes, exons, introns, promoters) and CpG islands. The minimum size for an overlap was 1 bp. The program g:Profiler was used for functional enrichment analyses of overlapping genes with DMRs.

### 2.8 RNA-Seq library construction and data analysis

One whole quadriceps muscle per mouse (N=4 mice/group) was subjected to bulk RNA sequencing (Medgenome). Total RNA was processed for library construction (**Supplementary Methods 1.3)**. For analyses, 32 pair-end raw sequence reads files (fastq format) were transferred to Illumina Partek Flow server producing 16 paired RNA-seq samples. Analysis was performed using Illumina Partek Flow RNA-Seq pipeline (**Supplementary Methods 1.3**). Volcano plots were generated highlighting the top 50 differentially regulated genes (i.e., greatest absolute fold changes). Gene Set Enrichment Analyses (GSEA) were generated and hierarchically clustered with DESeq2 using an R bioconductor package (v3.14.3). Pathways with enrichment score of ≥ 3.0 and p-value < 0.05 were considered significant. For differential analysis of the RNA-seq data, normalized gene counts were analyzed via 2×2 ANOVA (“treatment” and “genotype” as the factors) in Illumina Partek Flow.

### 2.8 Conjoint analysis

Conjoint analysis was performed to integrate the DNA methylation and RNA sequencing datasets (with matched columns) and identify overlapping genes with a high correlation across different comparison groups using Biomni tool, as recently described [16].

### 2.9 Statistics

In addition to analyses already described for methylation and RNAseq studies, experiments comparing two groups were analyzed with Student’s t-tests, and experiments comparing ART effects across WT and Tg26 animals were analyzed via two-way ANOVA (factor 1=treatment: FTC or VEH; factor 2=Genotype: WT or Tg26) with interaction, followed by post-hoc testing with Fisher’s Least Significant Difference tests where appropriate. For qPCR datasets, statistical analyses were performed on delta delta CT values, and relative fold changes were graphically represented. All analyses were performed with JMP Pro v.18; p-values of p<0.05 were defined as significant and p-values 0.05≤ p≤ 0.1 were noted as trends. Data from vivo studies are represented in box plots as median, quartiles, and outlier fences (where available; first quartile - 1.5* (interquartile range) and third quartile + 1.5*(interquartile range)); each data point shown represents one mouse.

## 3. RESULTS

### 3.1 Short-term FTC treatment did not affect body composition nor induce gross deficits in skeletal muscle but affected myogenetic differentiation in Tg26 mice

As ART-induced muscle dysfunction develops gradually in PLWH [17], we did not anticipate functional deficits following a short, 1 month course of ART. Consistent with this idea, neither hang time (p_treatment_ = 0.4403; **Figure 1A**) nor hang time normalized to body weight (p_treatment_ **=** 0.5518; **Figure 1B**) showed an impact of ART. Hang time was not different between WT and Tg26 mice immediately prior to sacrifice (p_genotype_ = 0.6808), nor were interactions between genotype and ART administration observed (p_interaction_ = 0.1532). Similarly, FTC did not cause longitudinal changes in hang time over the study (i.e., in terms of percent change from baseline, p_treatment_ = 0.9144; **Figure 1C**), although Tg26 mice demonstrated a 2-fold greater decline in hang time over the study as compared to WT mice (p_genotype_ = 0.0453, **Figure 1C**). Consistent with lack of changes in muscle function, no differences were observed in muscle fiber size (**Figure 1D-E**) or minimum Feret diameter (**Figure 1F**) across genotypes or treatments. While longitudinal changes in body composition (%lean or %fat; **Figure 2A-B)** were not affected by genotype (p>0.2403) or FTC (p>0.3530) in this short-term study, a trend was observed for %fat, which tended to be selectively increased by FTC only in the Tg26 mice over the study (p=0.0854, **Figure 2B**).

**Figure 1:**
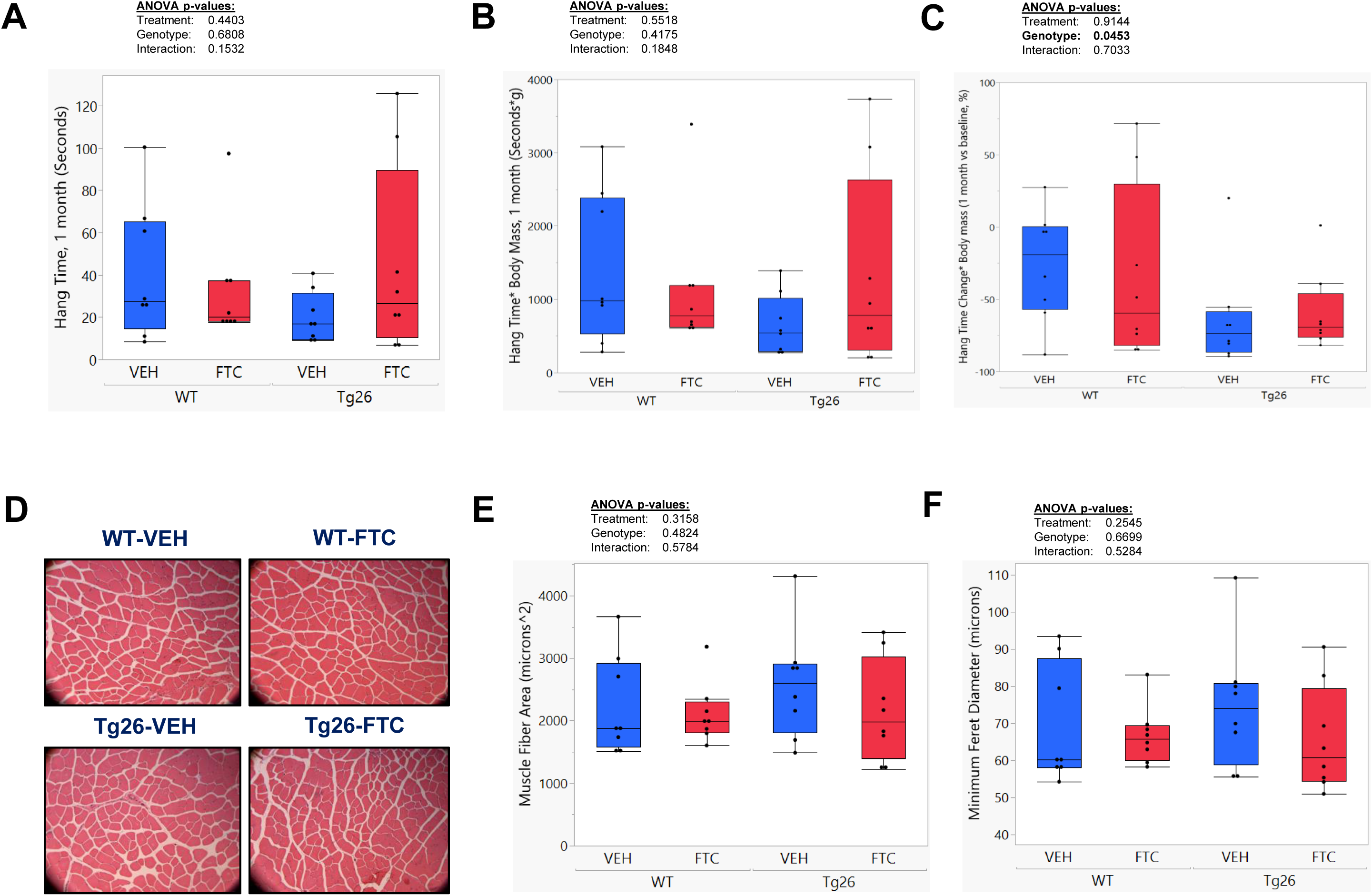
Short-term FTC treatment induced few, if any, functional or structural changes in skeletal muscle. Neither hang time (A) nor hang time normalized to body mass (B) was affected by the HIV genotype background or FTC treatment. While percent change in hang time as compared to baseline (C) was also unaffected by FTC treatment, this metric was significantly lower in the Tg26 HIV mice as compared to WT controls (p_genotype_ = 0.0453). (D) Representative cross-sectional H & E-stained images of quadriceps muscles from each group and quantitative analyses demonstrated that neither mean muscle fiber area (E) nor minimum ferret diameter (F) were affected by genotype or FTC treatment. Box plots show median, quartiles, and outlier fences for each group, where outlier fences represent first quartile −1.5* (interquartile range) and third quartile +1.5* (interquartile range), and each black circle represents one mouse. P-values from two-way ANOVA analyses comparing groups are shown above each graph.

**Figure 2:**
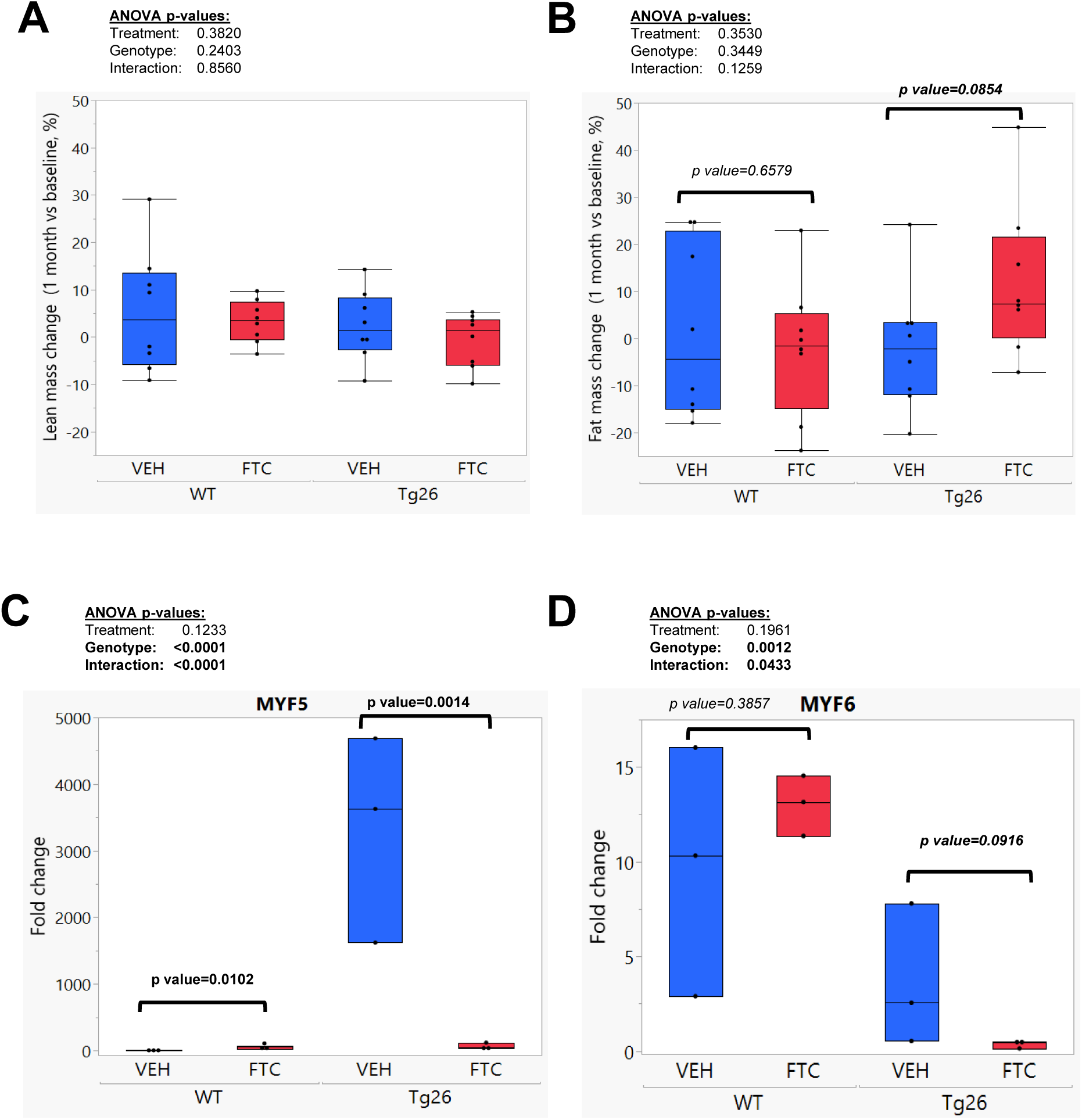
Short-term FTC treatment induced few, if any, functional or structural changes in skeletal muscle. DXA analyses showed no effect of genotype or treatment on changes in longitudinal measurements of lean mass (A) or fat mass (B) although pairwise comparisons showed a trend for increased changes in fat mass in the FTC-treated Tg26 mice (p=0.0854). (C) Gene expression levels of the immature myogenic marker Myf5 were greater in quadriceps muscles of Tg26 mice (VEH-treated) compared with WT-VEH mice and were interestingly increased by FTC in the WT mice but further decreased by FTC in the Tg26 mice (p_interaction_ <0.0001). (D) Gene expression levels of the mature myogenic marker Myf6 were lower in the Tg26- as compared to WT mice and tended to be further suppressed by FTC treatment in the Tg26 mice (p_interaction_ = 0.0433). Box plots show median, quartiles, and outlier fences (where available) for each group, where outlier fences represent first quartile −1.5* (interquartile range) and third quartile +1.5* (interquartile range), and each black circle represents one mouse. P-values from two-way ANOVA comparing groups are shown above each graph; groups with different superscript letters are significantly (p<0.05) different from one another as shown by post-hoc pairwise comparisons.

Considering the lack of structural and functional changes between groups, we quantified the mRNA expression of early (*Myf5)* and late *(Myf6)* myogenic regulators to determine whether markers of cell maturation were impacted by HIV phenotypes and/or ART administration. Interestingly, an interaction (p_interaction_ <0.0001) was observed for *Myf5*, where *Myf5* was mildly but significantly increased by FTC in WT (p=0.0102) but downregulated by FTC in Tg26 mice (p=0.0014), suggesting a uniquely detrimental effect of FTC in the HIV background **(Figure 2C)**. An interaction (p_interaction_ = 0.0433) was also observed for *Myf6*, where *Myf6* was unchanged by FTC in WT (p=0.3857) but tended to be downregulated by FTC in Tg26 mice (p=0.0916; **(Figure 2D**). This suggested that myogenic cell maturation was disrupted in the HIV model as compared to WT littermates, and that FTC exacerbated this trend, consistent with clinical reports of muscle dysfunction in PLWH on long-term ART [18]. This supports the appropriateness of the FTC administration model used here for investigating molecular mechanisms upstream of the gradual decline in muscle function seen in PLWH on ART, prompting further investigation into the impact of HIV phenotypes and ART on the skeletal muscle DNA methylation landscape and transcriptome.

### 3.2 An HIV phenotype (Tg26) and FTC treatment induced altered methylation patterns in genes relevant to skeletal muscle function

Methylation results were considered in three separate contexts of biological relevance: **1.** Impact of HIV phenotype (i.e., Tg26-VEH vs. WT-VEH, showing HIV phenotype effects independent of ART and mimicking the pathogenesis of untreated HIV), **2.** Impact of ART treatment (i.e., comparing WT-FTC to WT-VEH, showing FTC effects independent of altered HIV biology), and **3.** Impact of ART in HIV (i.e., comparing Tg26-FTC to Tg26-VEH, showing effects of ART in the context of HIV biology, mimicking the pathogenesis of treated HIV). Although not discussed in detail here due to lower translational relevance, we also considered comparisons between ART-treated WT vs Tg26 mice (i.e., Tg26-FTC vs. WT-FTC) to balance the experimental design and reveal whether the impact of HIV biology (i.e., Tg26 vs. WT) on the muscle epigenetic landscape was impacted by ART (**Supplementary Results 1**).

#### 3.2.1 : Impact of an HIV phenotype on DNA methylation patterns

The effect of an HIV phenotype on methylation patterns in skeletal muscle was assessed between VEH-treated WT and Tg26 mice, revealing 19 DMRs as hypermethylated and 15 DMRs as hypomethylated in Tg26-VEH vs. WT-VEH mice (**Figure 3A**). Among the 151 significant DMCs detected, 70 were hypermethylated and 81 were hypomethylated in the Tg26-VEH vs. WT-VEH mice (**Figure 4A**).

**Figure 3:**
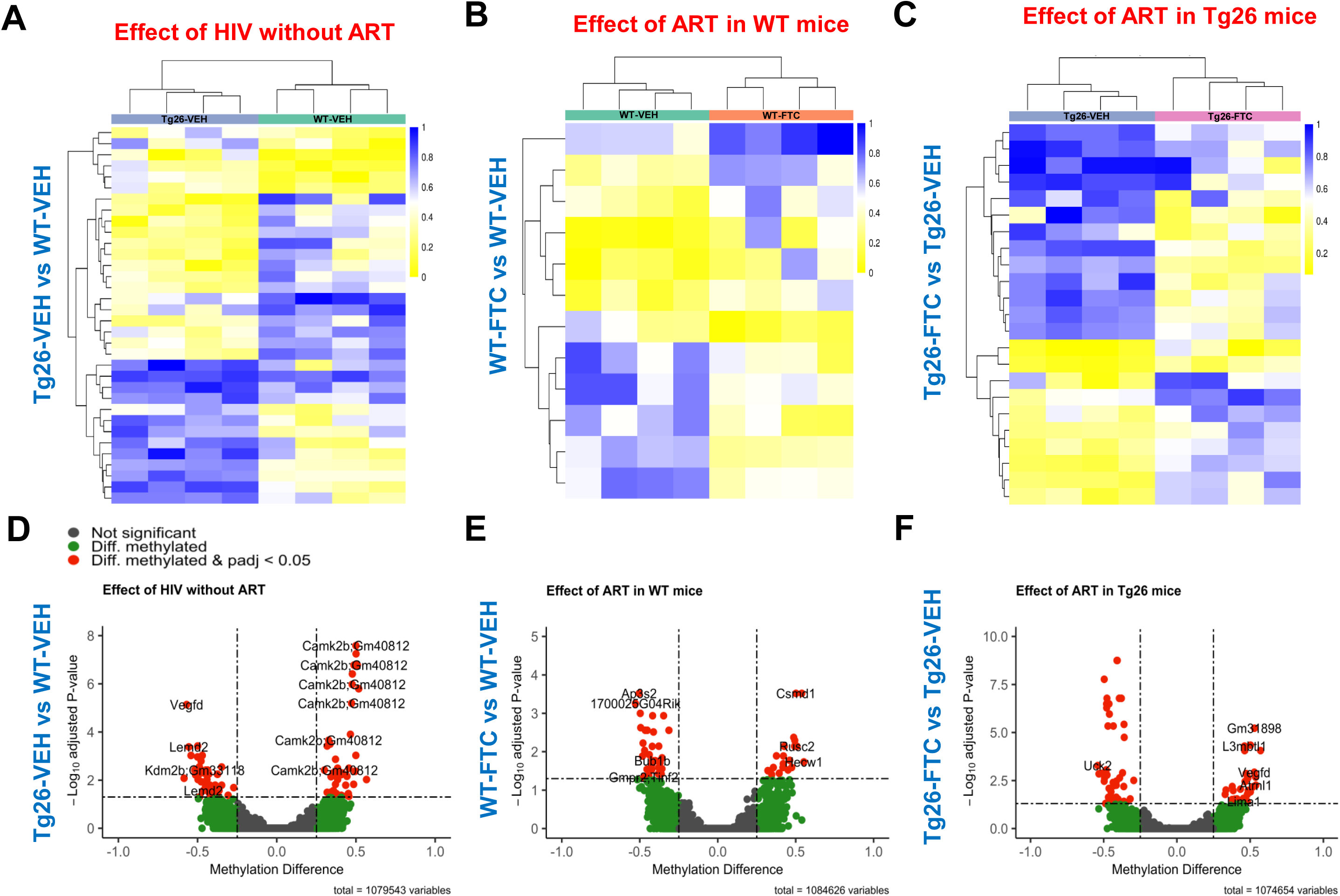
Impact of HIV phenotype and ART on WT and Tg26 mice skeletal muscle DNA methylation patterns. (A-C): Differentially methylated regions (DMRs) between the comparison groups: (A) Tg26-VEH vs WT-VEH, (B) WT-FTC vs WT-VEH and (C) Tg26-FTC vs Tg26-VEH. Heatmaps of DMRs identified using bisulfite sequencing methylation analysis are shown. DMRs are represented as rows, and each column represents one sample. The color intensity reflects from yellow to blue as degree of methylation in each group (ranging from 0, or 0% methylated, to 1, or 100% methylated). (D-F): Genes associated with differentially methylated CpG sites (DMCs) represented in volcano plots are shown for the following comparison groups: (D) Tg26-VEH vs WT-VEH (E) WT-FTC vs WT-VEH and (F) Tg26-FTC vs TG26-VEH.

**Figure 4:**
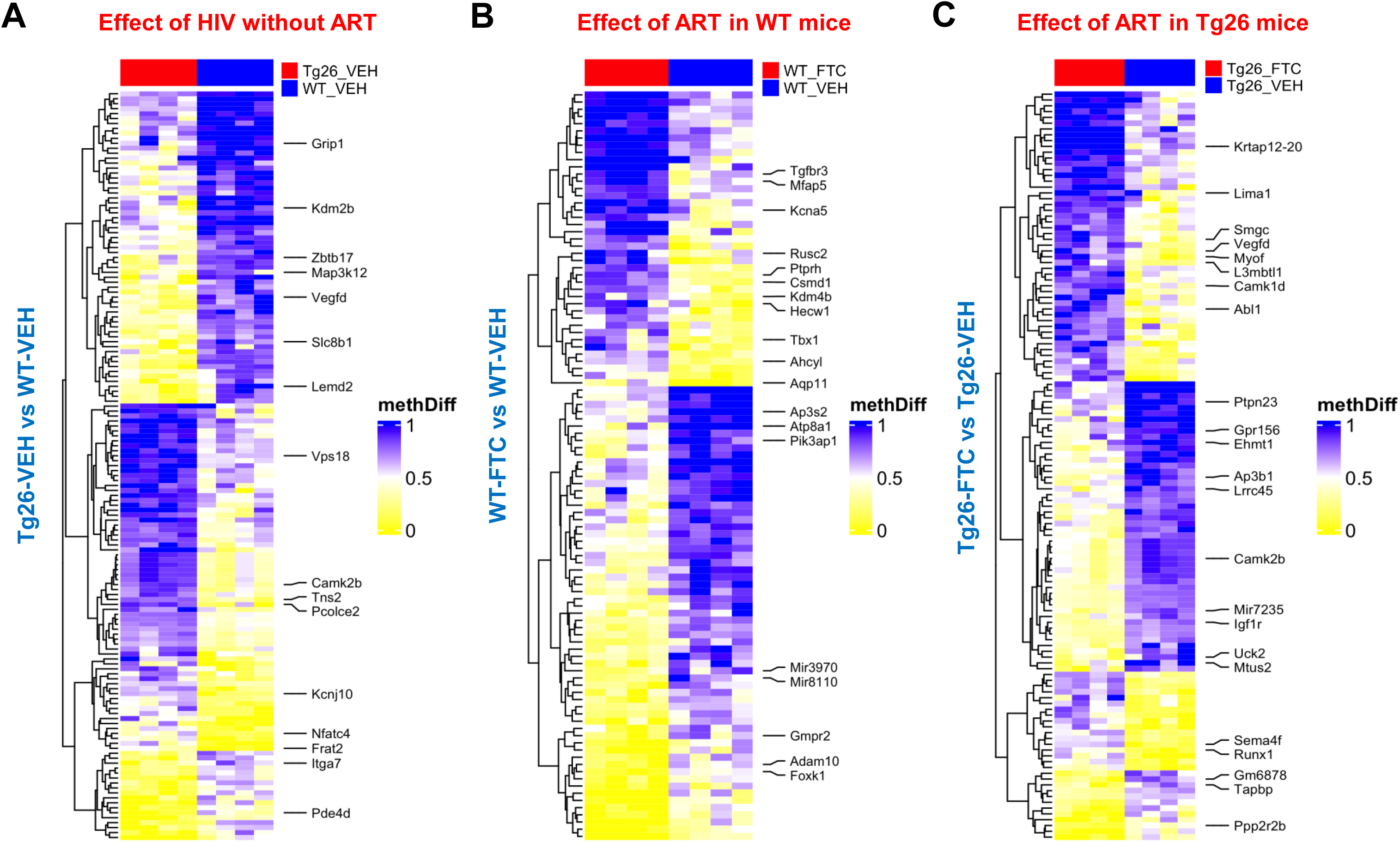
Heatmaps of differentially methylated CpG sites (DMCs) in skeletal muscle tissue across the following experimental groups: Tg26-VEH vs WT-VEH (A), WT-FTC vs WT-VEH (B), Tg26-FTC vs Tg26-VEH and (C). DMCs were identified using bisulfite sequencing methylation analysis method and are represented as rows, and each column represents one sample. Values are indicated as methylation difference (methdiff) between groups ranging from 0 (no difference) to 1 (maximum difference). Color intensity denotes the degree of methylation difference. Both DMCs and samples were clustered to highlight the patterns of methylation variation across the samples.

Only one pathway, “*Midface Retrusion*”, was associated with the significant DMRs (**Supplementary Table 1**). *Camk2B* showed the highest hypermethylation score in Tg26-VEH vs. WT-VEH mice at three locations which contained 12, 8 and 4 CpGs (Chr11: 6006422 – 6006618; 197 bp; 6007141 – 6007240; 100 bp; 6007943 −6008072; 130 bp). *Pcolce2* (Chr9: 95638732 – 95639031; 300 bp) with 5 CpGs was the second most hypermethylated gene. Additionally, among the genes hypomethylated in Tg26-VEH mice, *Lemd2* (Chr17: 27194732 – 27194783; 52 bp) with 5 CpG’s had the highest score (**Supplementary Table 2, Figure 3D**).

#### 3.2.2 : Impact of ART alone on DNA methylation patterns

Comparing WT mice treated with and without ART (i.e., WT-FTC vs. WT-VEH, ART effects without confounding effects of disease) identified 6 DMRs as hypermethylated and 6 as hypomethylated in WT-FTC vs. WT-VEH (**Figure 3B**). Regarding DMCs, 41 sites were hypermethylated and 63 sites were hypomethylated in WT-FTC vs. WT-VEH (**Figure 4B**).

Genes associated with differential methylation were related to cellular components and molecular function, with the pathway “*Long Term Depression”* enriched by ART treatment (**Supplementary Table 1**). Genes within the DMRs such as *Gnas* (chr2: 174284865 – 174285126; 262 bp length, 27 CpG sites) and *Myom1* (chr17: 71022014 – 71022322; 309 bp length, 4 CpG sites), were hypermethylated with FTC treatment, whereas genes including *Gdpd5* (Chr7: 99383798 – 99384074; 277 bp length, 4 CpG sites) and *Timp2* (Chr11: 118343438 – 18343533; 96 bp length with 4 CpG sites) demonstrated hypomethylation with FTC (**Supplementary Table 2, Figure 3E**).

#### 3.2.3 : Impact of ART in the HIV background on DNA methylation patterns

The effect of ART in an HIV background (i.e., Tg26-FTC vs. Tg26-VEH, mimicking the pathogenesis of HIV treatment) revealed 10 hypermethylated DMRs and 13 hypomethylated DMRs in the Tg26-FTC vs. Tg26-VEH mice (**Figure 3C**), where 62 DMCs were hypomethylated and 67 hypermethylated with FTC (**Figure 4C**). Twelve pathways were identified from these DMRs, with most genes associated with the “*Comprehensive Resource of Mammalian Protein Complexes (CORUM)*” pathway informing protein-protein interactions and protein complexes (**Supplementary Table 1**). Among the 10 hypermethylated DMRs, *Lima1* (Chr15: 99803941 – 99804234; 294 bp length, 7 CpG sites) and *L3mbtl1* (Chr2: 162948216 – 162948523, 308 bp length, with 11 CpG sites) exhibited the highest methylation scores in the Tg26-FTC mice. One gene, *Camk2B,* exhibited hypomethylation in the Tg26-FTC mice at two distinct locations (Chr11: 6006422 – 6006618; 197 bp length and 6007141 – 6007240; 100 bp length) (**Supplementary Table 2, Figure 3F)**.

Taken together, the differential methylation patterns and pathway enrichment analyses suggest that presence of an HIV phenotype (Tg26) and exposure to an ART (FTC) altered methylation of skeletal muscle genes in a manner consistent with eventual functional impairment.

### 3.3 RNAseq revealed FTC administration promoted differential gene expression across WT and Tg26 mice

RNAseq comparisons of differentially expressed genes (DEG) were made in the same three biological contexts as methylation analyses, with a 4^th^ consideration of ART-treated WT and Tg26 mice (i.e., Tg26-FTC vs. WT-FTC) considered to balance the experimental design shown as a supplementary analysis (**Supplementary Results 2, Supplementary Figure 3A-C**). We also added a 2×2 ANOVA design, permitting consideration of the general impact of drug treatment and the general impact of HIV background (**Supplementary Results 3.1-3.2, Supplementary Figure 4A-F**).

#### 3.3.1 : Impact of an HIV phenotype on skeletal muscle transcriptome

Comparing Tg26-VEH vs WT-VEH identified 744 (380 protein coding) DEG, with 458 genes downregulated and 286 genes upregulated in the Tg26-VEH as compared to WT-VEH mice (**Figure 5A**). Gene Set Enrichment Analysis (GSEA) identified 28 enriched pathways, with the *“Neuroactive ligand-receptor interaction*” pathway showing the highest enrichment score (**Supplementary Table 3**), where genes associated with the gene ontology category “*cellular anatomical entity*” and regulation of biological processes (**Figure 5B**). Genes such as *LEP*, *ADORA1*, *ADIPOQ*, *FFAR2* and *REX2* were downregulated, while *IGFLR1*, *ALDH1B1*, *PPARα*, *WNT5b* and *MYH1* were upregulated in the Tg26-VEH mice (**Supplementary Figure 2A**).

**Figure 5:**
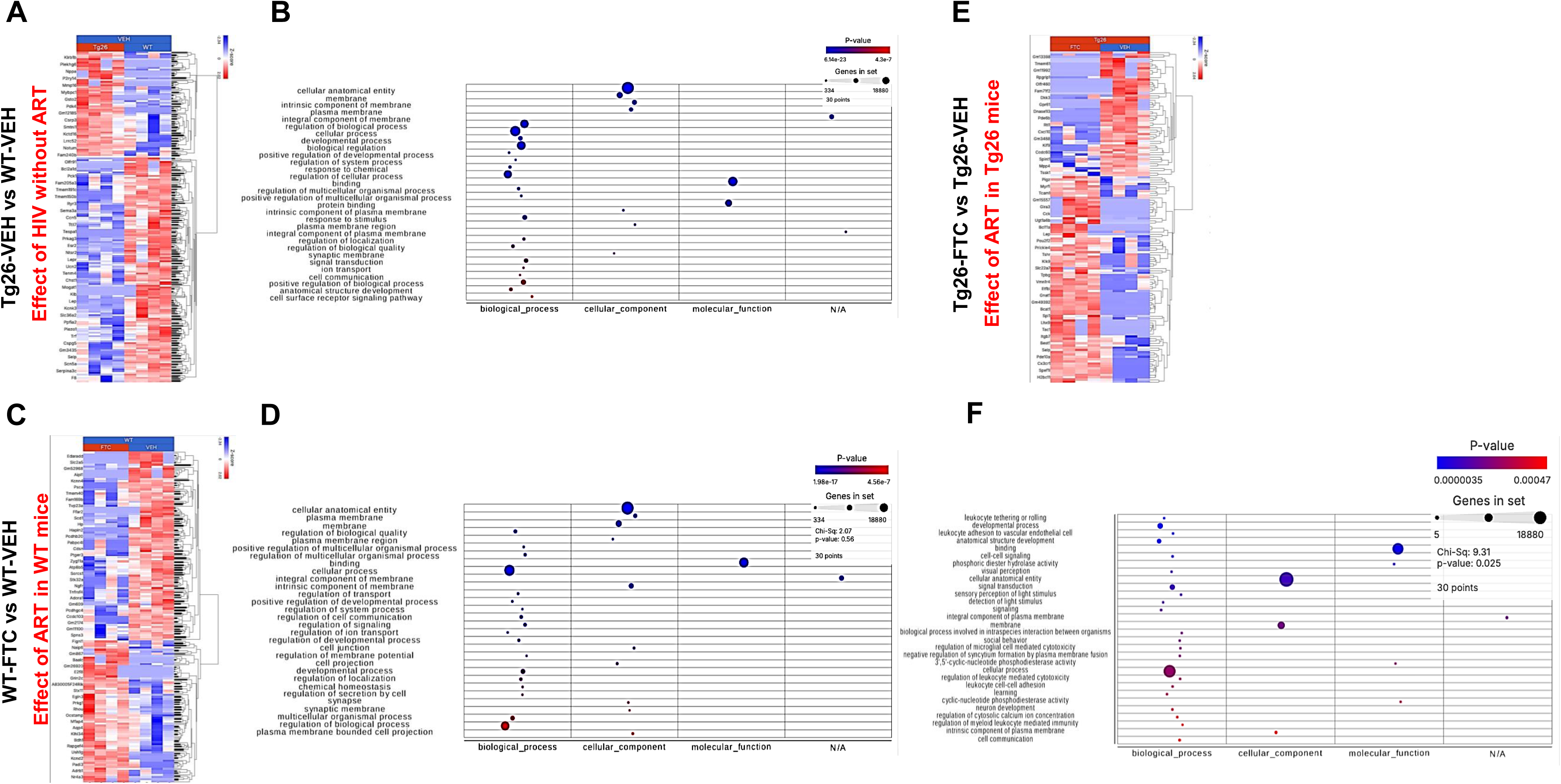
Impact of HIV phenotype and ART on WT and Tg26 mice skeletal muscle transcriptome. Comparison across the groups: Tg26-VEH vs WT-VEH (A&B), WT-FTC vs WT-VEH (C&D), and Tg26-FTC vs Tg26-VEH (E&F). (A, C, E): Heatmap shows genes (rows) and individual mice from each group (columns), where color intensity reflects the Z-score, with blue indicating downregulation and red indicating upregulation. (B, D, F): Gene Ontology plots of protein coding genes and biological processes compared between Tg26-VEH vs WT-VEH (B), WT-FTC vs WT-VEH (D), and Tg26-FTC vs. Tg26-VEH (F).

#### 3.3.2 : Impact of ART alone on skeletal muscle transcriptome

WT mice treated with ART (i.e., WT-FTC vs WT-VEH) demonstrated 659 DEG (322 protein coding) with 265 upregulated and 394 downregulated by FTC (**Figure 5C**). Pathway enrichment analysis revealed “*Neuroactive ligand-receptor interaction”* and *“Regulation of lipolysis in adipocytes”* associated with the DEG (**Supplementary Table 3**), and most of the DEG belonged to *“cellular component”* or *“cellular biological process”* gene ontology categories (**Figure 5D**). ART downregulated genes including *HPCA*, *ADORA1, ADIPOQ, LEP, LPAR4, TSHR, KMO*, while expression of *ADRB1, IL4I1, CCK, PPAR-α, ALDH1B1* and *PRKG1* were upregulated by FTC (**Supplementary Figure 2B**).

#### 3.3.3 : Impact of ART in the HIV background on skeletal muscle transcriptome

Analysis of ART treatment in the HIV background (Tg26-FTC vs Tg26-VEH) revealed 375 DEG (146 protein coding) with 170 upregulated and 205 downregulated by FTC treatment (**Figure 5E**). A total of 12 enriched pathways were observed, with *“Neuroactive ligand-receptor interaction”* representing the largest enrichment score (**Supplementary Table 3**). Most DEG associated with gene ontology categories of *“cellular anatomical entity”* and regulation of biological processes (**Figure 5F**). Genes including *AVPR1A* and *GABRB3* associated with *“Neuroactive ligand-receptor interaction”* were downregulated whereas genes including *LEP*, *CCK*, *ADRA1*, *LPAR4*, *TSHR*, *NPY6R* and *TAC1* were upregulated by FTC (**Supplementary Figure 2C**).

Collectively, the DEG profiles and corresponding enriched pathway analyses further support the idea an HIV phenotype (Tg26) and ART treatment (FTC) impacted skeletal muscle gene regulation via key pathways including cellular differentiation, regulation of lipid metabolism, neuroactive ligand-receptor interactions, cytokine interactions, cell adhesion interactions, the Wnt signaling pathway, oxidative stress, and genes related to regulation of muscle strength.

### 3.4 Conjoint analysis revealed minimal functional overlap between transcriptome and methylome datasets across contexts of biological relevance

Conjoint analysis of genes associated with sites of differential methylation (i.e., DMC) and DEG from RNA seq data revealed 3 overlapping genes in the comparison group assessing impact of an HIV phenotype (Tg26-VEH vs WT-VEH). These included *MID-1, Gpr156* and *D830013O20Rik*, showing a strong co-relation in both datasets (**Supplementary Figure 5A, E)**. This supports the existence of coordinated epigenetic and transcriptomic changes attributable to an HIV phenotype, with evidence for transcriptional activation patterns. No direct gene-level overlap was found between the methylome and transcriptome datasets in the comparison addressing ART effects without confounding effects of disease (i.e., WT-FTC vs WT-VEH; **Supplementary Figure 5B, F**), nor the comparison facilitating analysis of ART treatment in the HIV background (Tg26-FTC vs Tg26-VEH) (**Supplementary Figure 5C, G)**. However, conjoint analyses revealed one overlapping gene (*TMEM266*) in the comparison between ART-treated WT vs Tg26 mice (i.e., Tg26-FTC vs. WT-FTC) (**Supplementary Figure 5D, H)**.

## 4. DISCUSSION

DNA methylation is an important epigenetic mechanism by which skeletal muscle cells regulate transcriptional expression, such as for the regulation of protein-coding genes responsible for sarcomere organization [19]. While it is well established that HIV infection and usage of ART drugs associates with skeletal muscle fatigue and weakness [20], the underlying mechanisms associated with these changes in muscle are still largely undefined. To the best of our knowledge, results presented here represent the first study to assess how an HIV phenotype and usage of an ART may impact skeletal muscle DNA methylation and transcriptome patterns. Our data show that the combined presence of HIV viral proteins (i.e., the Tg26 model) and administration of an ART (FTC) promoted striking alterations in DNA methylation patterns and associated changes in gene transcriptional patterns.

It is important to note that the methylation and transcription-related changes observed here occurred largely in the absence of skeletal muscle functional deficits. As mentioned previously, since ART-induced muscle dysfunction develops gradually in PLWH [17, 21], we did not anticipate observing functional performance deficits in our experimental design. We did, however, detect a reduction in muscle endurance in the Tg26 as compared to WT mice over the course of the study, consistent with previous reports of muscle atrophy in Tg26 as compared to WT controls [14]. The muscle-wasting phenotype of Tg26 mice is more profound in fast-twitch (glycolytic) fibers and appears first in fast-twitch muscles like the EDL [14]. The four-limb hang time test employed in our studies primarily tests the function of fast-twitch muscles involved in grip including the gastrocnemius, TA, and EDL [22], and therefore the reduction in hang time observed in Tg26 vs. WT mice in our study may be consistent with a reduction in glycolytic fibers. In addition, although we did not detect functional deficits from FTC in these short-term studies, we did observe extraordinarily high expression of the early myogenic regulatory gene *Myf5* in the Tg26 mice that was uniquely reduced by FTC treatment, and a decrease in the later myogenic regulatory gene *Myf6* that tended to be more substantial in the FTC-treated Tg26 mice. This raises the possibility that myoblast differentiation and maturation patterns may be arrested in the context of an HIV phenotype and ART administration. This outcome is consistent with the transcriptome- and methylome-related changes observed in the bioinformatics studies suggesting that the combination of an HIV phenotype and exposure to FTC treatment induced regulatory changes positioned to be detrimental to skeletal muscle. Longer-term treatment studies will be needed to determine if the experimental conditions here would result in functional impairment given a sufficient duration of treatment.

DMRs associated with exposure to FTC in the absence of a disease phenotype (i.e., WT-FTC vs. WT-VEH) were enriched in biological processes related to the pathway “*Long term depression.”* Interestingly, many of the hypermethylated genes such as *Myom1*, *Gnas* and *Tsks* participate in processes related to skeletal muscle integrity, regeneration and myofiber development [23–26], suggesting that hypermethylation of these genes, and the subsequently expected transcriptional repression, would be detrimental to skeletal muscle. Similarly, DMRs associated with presence of an HIV phenotype in absence of treatment (i.e., Tg26-VEH vs. WT-VEH) were related to the pathway “*Midface retrusion.”* While ‘midface retrusion’ describes a process by which the infraorbital and perialar regions of the face are underdeveloped (suggesting bony changes, rather than muscular), the genes associated with this process included *Camk2B*, which was hypermethylated in Tg26 mice. The *Camk2B* gene plays an important functional role in neurotransmitter synthesis and synaptic organization [27], glucose uptake in skeletal muscle, and cardiac function [28]. Hypermethylation of *Camk2B* may be consistent with impaired neuromuscular junction biology that could contribute to the skeletal muscle wasting observed in an HIV phenotype. Another hypermethylated gene, *Pcolce2,* is a regulator of lipid metabolism [29] and marker of fibro-adipogenic progenitor cells which are crucial to the support of muscle stem cells and to processes of muscle regeneration [30]. Consistent with this finding, RNAseq showed ‘*Regulation of lipolysis*’ and ‘*Adipocytokine signaling*’ as enriched pathways in Tg26-VEH as compared to WT-VEH mice, with key adipogenic genes like *ADIPOQ* and *LEP* showing lower expression in Tg26-VEH as compared to WT-VEH mice. Unfortunately, we did not detect mRNA-level changes of differentially methylated genes in RNAseq analyses; it is possible and perhaps even likely that the time course of transcriptional changes may be different than that of the epigenetic regulatory mechanisms predicted to affect their expression. Although beyond the scope of the current study, in vitro studies employing primary myoblast-derived myofibers would be an excellent tool by which to mechanistically investigate the time course and functional connections between these two biological processes induced by HIV proteins and exposure to ARTs.

It is intriguing that patterns of differential methylation induced by the presence of an HIV phenotype were quite different in the FTC treated mice (Tg26-FTC vs. WT-FTC) as compared to the VEH treated mice (Tg26-VEH vs. WT-VEH), where many of the genes found to be methylated in the ART-treated Tg26 mice also suggest a connection to muscle function. The hypermethylated *HOXA2* gene is required for muscle primordia development [31], and the hypermethylated *Mbnl2* gene is critical for skeletal muscle regeneration [32]; reduced expression of *Mbnl2* promotes myotonic dystrophy I [33]. In contrast, FTC treatment of Tg26 mice promoted hypomethylation of *Camta1* which plays a crucial role in several processes including embryonic development [34, 35]. Given the altered expression patterns, we observed for myogenic regulatory genes *Myf6* and *Myf5* with FTC treatment in Tg26 mice, it is possible that these changes in DNA methylation patterns of muscle development-related genes could tie to functional changes in myoblast maturation.

FTC responses within Tg26 mice, mimicking the biology of HIV treatment with ART, identified genes annotated to regions related to the Wnt signaling and transforming growth factor beta2 production pathways. The hypermethylated gene *Lima1*, a regulator of the actin cytoskeleton, is expressed in early stages of skeletal muscle development and plays an important role in muscle cell growth [36]. Hypermethylation of *Lima1* with ART in the Tg26 mice may be consistent with muscle fiber atrophy. Excitingly, we observed the *Camk2B* locus, which was hypermethylated in the Tg26-VEH as compared to WT-VEH mice (i.e., hypermethylated in an untreated HIV phenotype biological context), became hypomethylated following FTC treatment in the Tg26 mice. Thus, although the combined exposure to HIV proteins and ART drugs is predicted to be synergistically detrimental, mimicking the ART-induced muscle dysfunction that develops gradually in PLWH [17, 21], further work building upon these exploratory analyses will ultimately be needed to test the contributions of discrete genes to these mechanisms.

Previous RNAseq studies showed that adverse effects of ART may be linked to altered fat metabolism and fat redistribution [37, 38], and lipodystrophy is frequently found in in PLWH on ART, particularly with older ART regimens [39]. Numerous studies have investigated the correlation between serum leptin levels, virological load, and immune state in PLWH [40]. We observed suppressed expression of leptin (*LEP*) in skeletal muscle of Tg26-VEH as compared to WT-VEH mice, and with ART treatment in the WT-FTC as compared to WT-VEH mice. Leptin is anabolic for muscle (**Supplementary Reference 1, 2**), and is abundantly expressed in skeletal muscle tissue, and therefore these trends for modulation of leptin by an HIV phenotype and ART could be well positioned to contribute to muscle dysfunction. While HIV patients with lower CD4 counts have lower serum leptin levels (**Supplementary Reference 3**), consistent with the suppressed expression of *LEP* found in the Tg26-VEH as compared to WT-VEH mice, hyperleptinemia was found in 14% of the ART-treated HIV patients with lipohypertrophy (**Supplementary reference 4**). In this context, it is notable that we saw a trend for increased whole-body %fat via DXA in the Tg26-FTC as compared to Tg26-VEH mice, consistent with the fact that *LEP* expression was uniquely increased by FTC in the Tg26 but not WT mice. The decreased expression of *LEP* in Tg26-VEH as vs. WT-VEH mice was paralleled by decreased mRNA expression of *ADIPOQ* (adiponectin) in Tg26-VEH vs. WT-VEH mice. Adiponectin drives mitochondrial biogenesis, glucose uptake, and fatty acid oxidation in muscle cells, and knockdown of this gene is associated with impaired muscle metabolism (**Supplementary Reference 5**). The lipodystrophy phenotype found in PLWH also promotes altered glucose homeostasis in skeletal muscle (**Supplementary Reference 6**), and *PPARα*, a gene important for lipid and glucose metabolism in skeletal muscle (**Supplementary Reference 7**), was upregulated by the presence of HIV proteins (i.e., in the Tg26-VEH vs. WT-VEH mice). Mechanistic studies interrogating the connections between lipid metabolism, glucose homeostasis, and the interactions between HIV biology and ART responses are warranted based on these findings and may shed light on the metabolic consequences of HIV and ART usage that could drive the early onset of musculoskeletal fragility.

Conjoint analysis of genes associated with both DMCs and DEGs (i.e., identified as being differentially expressed in both methylome and transcriptome datasets) was expected to uncover coordinated regulatory changes attributable to an HIV phenotype and ART usage, however very few overlapping genes were recognized in these analyses. Moreover, the specific roles of genes highlighted by conjoint analysis (*MID*-*1*, *Gpr156*, *D830013O20Rik*, and *TMEM266*) as related to skeletal muscle homeostasis and function are still unclear and will require further study. The lack of correlation between our methylome and transcriptome datasets may suggest that methylation and transcriptional changes operate through distinct or more complex regulatory mechanisms. Alternatively, it is possible that the dynamic connection between differential DNA methylation and transcriptional regulation may have occurred in a timeframe that was missed by assessing molecular endpoints only after 4 weeks of treatments; mechanistic, in vitro studies may be required to better understand the timing and regulatory connection between differential DNA methylation and transcriptional changes in skeletal muscle attributable to HIV and ART usage.

Our study has several limitations. We used a single dosage and short-term time course for FTC treatment, limiting understanding of any potential temporal dynamics or long-term effects on skeletal muscle. Moreover, the functional significance and underlying molecular mechanisms of altered DNA methylation and transcriptome endpoints need to be further investigated to better understand the interplay between FTC, epigenetic regulation, and muscle function. Although an HIV phenotype is reported to primarily affect fast-twitch fibers as noted above, our analyses were confined to bulk tissue analyses of mixed fiber type muscles (quadriceps), which may have negatively impacted our ability to detect differences between groups. Also, it is important to consider that studies here utilized a non-infectious mouse model; although the Tg26 model phenocopies the onset of HIV-induced frailty in humans [14], additional studies using infectious HIV models and using more translationally relevant tools like human myogenic cell lines will be needed to understand the translational relevance of the epigenetic and transcriptome changes induced by FTC.

Results shown here provide valuable insights into the mechanisms by which a clinically relevant ART (FTC) may drive DNA methylation and transcriptome changes to skeletal muscle in the context of HIV biology. Even a short course of FTC treatment was capable of significantly altering DNA methylation patterns and gene expression profiles within skeletal muscle tissue. In the Tg26 model of HIV, these modifications were associated with genes related to muscle growth, development and plasticity, and transcriptomic changes suggested alterations in pathways related to lipid metabolic processes. These analyses suggest novel targets for understanding, and eventually abrogating, the harmful effects of long-term ART use in PLWH on skeletal muscle mass and function, with the ultimate goal of increasing healthspan in this patient population.

## ACKNOWLEDGEMENTS

Funding for these studies was provided by the NIH / NIAMS (R01 AR082307). The authors would like to acknowledge the Augusta University’s Electron Microscopy and Histology Core (RRID:SCR_026810) for assistance with histological specimen preparation, and the Augusta University Cell Imaging Core (RRID:SCR_026799) for assistance with specimen imaging.

## AUTHOR CONTRIBUTIONS: CRediT author statement

**Alok Tripathi**: Data curation, Formal analysis, Investigation, Methodology, Visualization, Writing – Original Draft Preparation, Writing – Review & Editing

**Shabiha Sultana**: Investigation, Writing – Review & Editing

**Husam Bensreti**: Investigation, Writing – Review & Editing.

**Jie Chen:** Data curation, Formal analysis, Investigation, Methodology, Resources, Software, Visualization, Writing – Review & Editing.

**Arindam Paul**: Methodology, Resources, Software, Visualization, Investigation, Writing –

Review & Editing.

**Huidong Shi**: Data curation, Formal analysis, Funding acquisition, Investigation, Methodology, Resources, Software, Visualization, Writing – Review & Editing.

**Apeksha Anand**: Methodology, Software, Investigation, Writing – Review & Editing.

**Bharati Mendhe**: Investigation, Writing – Review & Editing.

**Kanglun Yu**: Investigation, Writing – Review & Editing.

**William E. Long**: Investigation, Writing – Review & Editing.

**Rodger D. MacArthur**: Funding acquisition

**Eric J. Belin de Chantemele**: Funding acquisition, Methodology, Project administration, Resources, Writing – Review & Editing.

**Mark W. Hamrick:** Conceptualization, Funding acquisition, Methodology, Project administration, Resources, Supervision, Writing – Review & Editing.

**Meghan E. McGee-Lawrence:** Conceptualization, Formal analysis, Funding acquisition, Methodology, Project administration, Resources, Supervision, Visualization, Writing – Original Draft Preparation, Writing – Review & Editing.

## Data availability statement

The data that support the findings of this study are available from the corresponding author upon reasonable request.

## Supplementary Methods

### 1.1 RNA isolation and qRT-PCR analysis

The mRNA expression levels of myogenic genes were quantified by real-time PCR using SYBR green (ThermoFisher #A25780) on a Bio Rad CFX Connect PCR system using 37.5ng cDNA per reaction. Gene expression was quantified using the delta delta Ct analysis method. Primer sequences included *MYF5* (F-TGAGGGAACAGGTGGAGAAC, R-CTGTTCTTTCGGGACCAGAC), *MYF6* (F-GCTAAGGAAGGAGGAGCAAA, R-GAAGAAAGGCGCTGAAGACT), and *GAPDH* (F-GGGAAGCCCATCACCATC, R-GCCTCACCCCATTTGATGTT) as a housekeeping gene.

### 1.2 Reduced representation bisulfite sequencing (RRBS) library preparation and bioinformatics analysis

A starting input of 500 ng genomic DNA was digested with 30 units of methyltransferases (MspI restriction enzyme) from New England Biolabs. Fragments were ligated to pre-annealed adapters containing 5’-methyl-cytosine instead of cytosine according to the manufacturer’s specified guidelines. Adaptor-ligated fragments ≥50 bp in size were recovered using the DNA Clean & Concentrator™-5 system (Cat#: D4003). The fragments were then bisulfite-treated using the EZ DNA Methylation-Lightning™ Kit (Cat#: D5030). Preparative-scale PCR was performed, and the resulting products were purified with DNA Clean & Concentrator™-5 (Cat#: D4003). Library quality control was performed on the Agilent 4200 TapeStation. Libraries were sequenced on an Illumina NovaSeq X platform (150 bp PE reads).

For pre-processing and quality control, input sequencing reads were trimmed using TrimGalore. Raw FASTQ files were adapted and quality trimmed. Post-trimming quality control was performed using FastQC. Picard tools were used to calculate the library insert size distribution. To process RRBS data, the trimmed reads were aligned to an in-silico bisulfite converted reference mouse genome assembly using Bismark. Methylation ratios for each cytosine in CpG context were called using MethylDackel. The methylation level of each sampled cytosine was estimated as the number of reads reporting a C, divided by the total number of reads reporting a C or T. Read depths per cytosine in the genome as well as in different genomic regions (e.g. gene body, promoter, CpG island, etc. based on available annotations) were calculated and tabulated using custom scripts.

### 1.3 RNA-Seq library construction and Data Analysis

Total RNA was extracted using the Promega Maxwell RSC simplyRNA tissue kit (cat#AS1340). Samples were homogenized in lysis buffer using a bead mill homogenizer. The homogenized lysate was then loaded onto the Maxwell RSC cartridge along with the necessary reagents for automated RNA extraction. Paramagnetic particles were used to capture RNA, which was then washed to remove contaminants and eluted in nuclease-free water. The resulting RNA was quantified using a Qubit Flex Fluorometer (Invitrogen Q33327) and assessed for integrity using an Agilent TapeStation 4200. Libraries were generated using the Illumina Stranded mRNA Prep kit (cat#20040532), following the manufacturer’s protocol with 300 ng RNA input. Briefly, polyadenylated RNA molecules were purified from total RNA using oligo-dT magnetic beads, and the captured mRNA was fragmented using divalent cations under elevated temperature. First strand cDNA synthesis was performed using random hexamer primers and reverse transcriptase, with Actinomycin D included to improve strand specificity. Following first strand synthesis, the RNA template was removed, and second strand cDNA synthesis was carried out using DNA polymerase I and RNase H. Strand specificity was maintained by incorporating dUTP in place of dTTP during second strand synthesis, enabling subsequent degradation of the second strand during post-PCR processing. The 3’ ends of the cDNA fragments were adenylated to prevent self-ligation, and adapters were ligated to prepare the cDNA for hybridization onto a flow cell. The adapter-ligated cDNA was enriched through PCR amplification and purified. Final library quality and quantity were assessed using a Qubit Flex Fluorometer (Invitrogen Q33327) for concentration measurements and an Agilent TapeStation 4200 for size distribution analysis. The libraries were sequenced on an Illumina NovaSeq 6000 platform for 300 cycles according to the manufacturer’s instructions.

The total reads range from 18 + million to 33 + million per sample with an average QA/QC score of at least 35.37. After pre-alignment QA/QC check and base trimming the sequencing reads were aligned to mm39 genome using the STAR aligner. The alignment rate ranges from 82.32% to 85.83% per sample with post-alignment QA/QC score of at least 38.66. These aligned reads were then quantified to annotation model mm39-emsembl-release-104 using Partek E/M algorithm, followed by data normalization (upper quantile, offset, CPM, and log base two transformation) to generate sequence counts of unique mouse genes for downstream differential expression analysis.

## Supplementary Results

### 1. Differences between ART-treated Tg26 vs. WT mice in DNA methylation patterns

Comparisons between ART treated Tg26 and WT mice demonstrated 9 hypermethylated and 9 hypomethylated DMRs in the Tg26-FTC vs. WT-FTC mice (**Supplementary Figure 1A**) with 42 DMC hypomethylated and 36 DMC hypermethylated (**Supplementary Figure 1B**) in the Tg26 mice. Functional enrichment analysis showed that Krüppel-like factors (KLFs), a DNA binding transcriptional factor, was associated with pathways that regulate metabolism and cellular mechanisms (**Supplementary Table 1**). Genes including *Mbnl2* (Chr14: 120287270 – 120287327; 58 bp), and *Hoxa2* (Chr6: 52163572 – 52163742; 171 bp, 4 CpG sites) were hypermethylated in the Tg26-FTC as compared to WT-FTC mice (**Supplementary Table 2, Supplementary Figure 1C**).

### 2. Differences between ART-treated Tg26 vs. WT mice in skeletal muscle transcriptome

Comparisons between ART treated Tg26 and WT mice revealed 463 DEG (173 protein coding), with 254 downregulated and 209 upregulated in the Tg26 mice (**Supplementary Figure 3A**). Pathways including ‘*Cell adhesion molecules’* and *‘Regulation of actin cytoskeleton’* were linked with the DEG (**Supplementary Table 3)**. The DEG were primarily associated with the *“cellular component’* and *“molecular function”* gene ontology categories (**Supplementary Figure 3B**). Genes such as *ITGAD, CDH2 CCK, LPAR5, NRXN1* were downregulated and genes including *CD4, CCL2, NRXN3, IL27 and DIAPH3* were upregulated in the FTC-treated Tg26 as compared to FTC-treated WT mice (**Supplementary Figure 3C**).

### 3. RNAseq analyses revealed generalized impacts of FTC administration across WT and Tg26 mice, and generalized impacts of an HIV phenotype across FTC and VEH-treated mice

#### 3.1 : Impact of drug treatment (FTC vs. VEH effects in genotypes combined)

Assessing the general impact of FTC across genotypes revealed 389 DEG (144 protein coding) with 231 downregulated and 158 upregulated in FTC vs. VEH-treated mice (**Supplementary Figure 4A**). Pathway enrichment analysis revealed *“Phototransduction”* and *“Neuroactive ligand-receptor interaction”* as the most highly enriched pathways (**Supplementary Table 3**). A majority of the DEG with FTC treatment belonged to the *“cellular anatomical entity”* and regulation of cellular processes gene ontology categories (**Supplementary Figure 4B**). Genes including *SLC2a5*, *LYPD6B*, *IL12Rβ2*, and *GDF7* were downregulated by FTC treatment, while *NPY6R ADRB1, CCK* were upregulated (**Supplementary Figure 4C**).

#### 3.1 : Impact of HIV background (Tg26 vs. WT effects in drug treatment groups combined)

Comparing across both drug treatment groups, 443 DEG (192 protein-coding) were detected, with 192 upregulated, and 251 downregulated in the Tg26 vs. WT mice (**Supplementary Figure 4D**). A total of 9 significantly enriched pathways were observed, with the *“Neuroactive ligand-receptor interaction”* demonstrating the highest enrichment score (**Supplementary Table 3**). DEG associated with the *“cellular component”* and *“biological process”* gene ontology categories (**Supplementary Figure 4E**). Genes including *ADRA2a*, *LEP*, *LEPR, CCK, ALDH3B2, NRXN-1 and CCL8* were downregulated and *CCL2, LDHAL6B, TIMP-1, and NRXN-3* were upregulated in the Tg26 as compared to WT mice (**Supplementary Figure 4F**).

## Supplemental Figure Captions

**Supplementary Figure 1:**
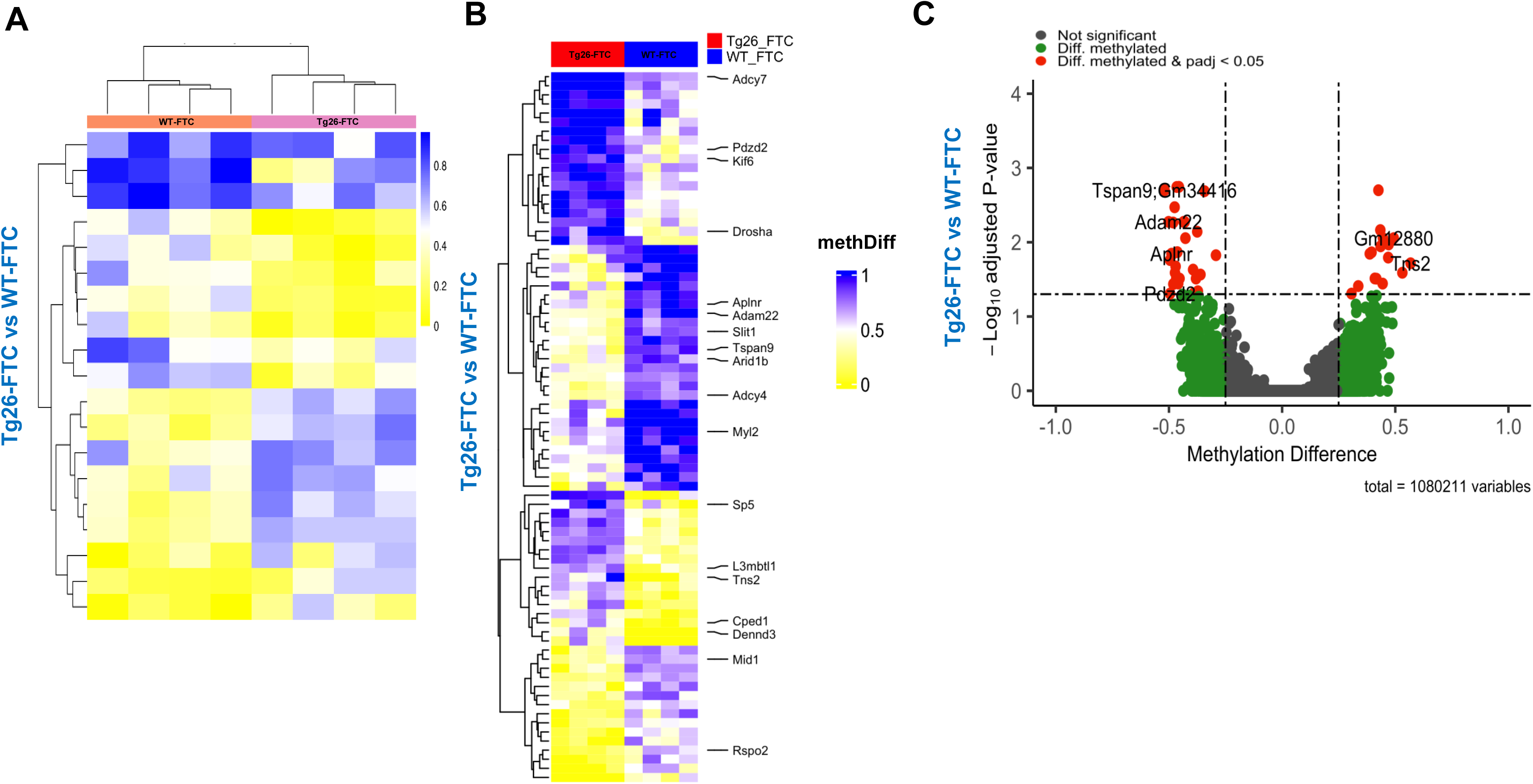
Impact of ART on WT and Tg26 mice skeletal muscle DNA methylation patterns: Differentially methylated regions (DMRs) between Tg26-FTC and WT-FTC mice groups: (A) Heatmap of DMRs identified using bisulfite sequencing methylation analysis. DMRs are represented as rows, and each column represents one sample. The color intensity reflects from yellow to blue as degree of methylation in each group. (B) Heatmap of DMCs was identified using bisulfite sequencing methylation analysis method and are represented as rows, while the experimental groups are represented as columns. Values are indicated as methylation difference (methdiff) between groups ranging from 0 (no difference) to 1 (maximum difference). Color intensity denotes the degree of methylation difference. Both DMCs and samples were clustered to highlight the patterns of methylation variation across the samples. (C) Genes associated with differentially methylated CpG sites (DMCs) represented in a volcano plot.

**Supplementary Figure 2:**
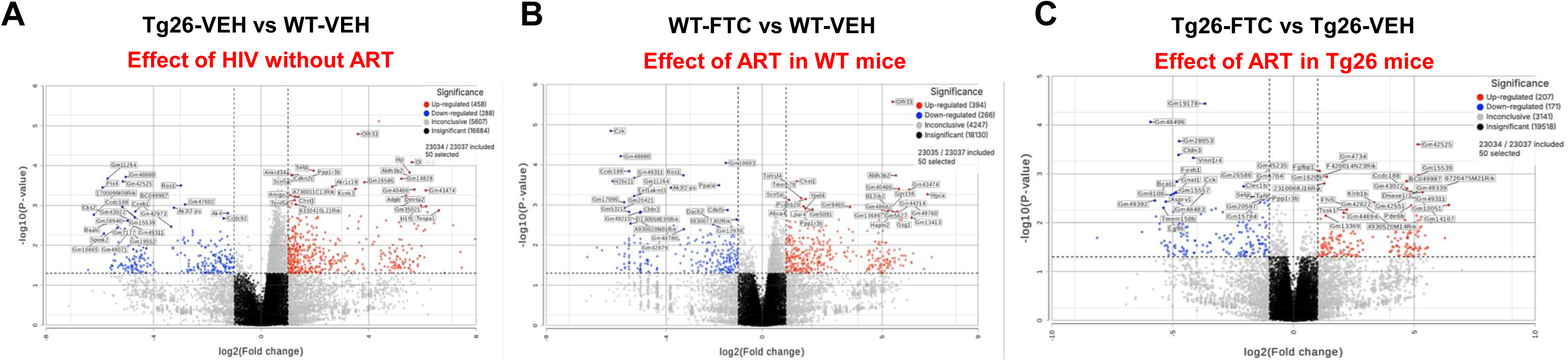
Impact of HIV phenotype and ART on WT and Tg26 mice skeletal muscle transcriptome: Volcano Plots. Volcano plots highlight the top 50 differentially expressed (upregulated and downregulated) genes among the comparison groups (**A**) Tg26-VEH vs WT-VEH, **(B**) WT-FTC vs WT-VEH, (**C)** Tg26-FTC vs. Tg26-VEH, based on log2 fold changes.

**Supplementary Figure 3:**
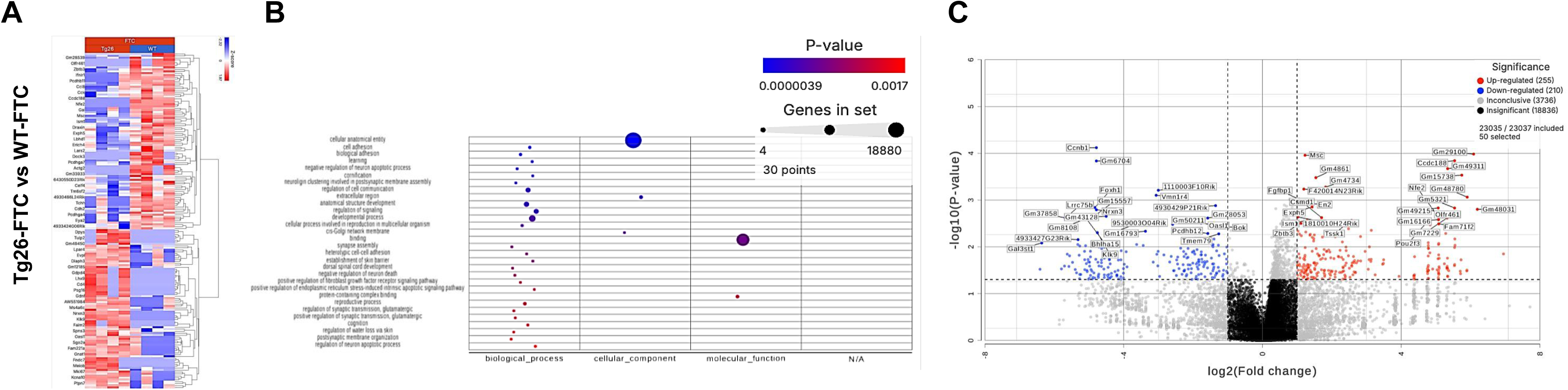
Impact of ART on WT and Tg26 mice skeletal muscle transcriptome: Comparison across Tg26-FTC vs WT-FTC mice. (**A**) Heatmap shows genes (rows) and individual mice from each group (columns), where color intensity reflects the Z-score, with blue indicating downregulation and red indicating upregulation (**B**) Gene Ontology plots of protein coding genes and biological processes compared between Tg26-FTC vs WT-FTC. (**C**) Volcano plots highlight the top 50 differentially expressed (upregulated and downregulated) genes between Tg26-FTC vs. WT-FTC based on log2 fold changes..

**Supplementary Figure 4:**
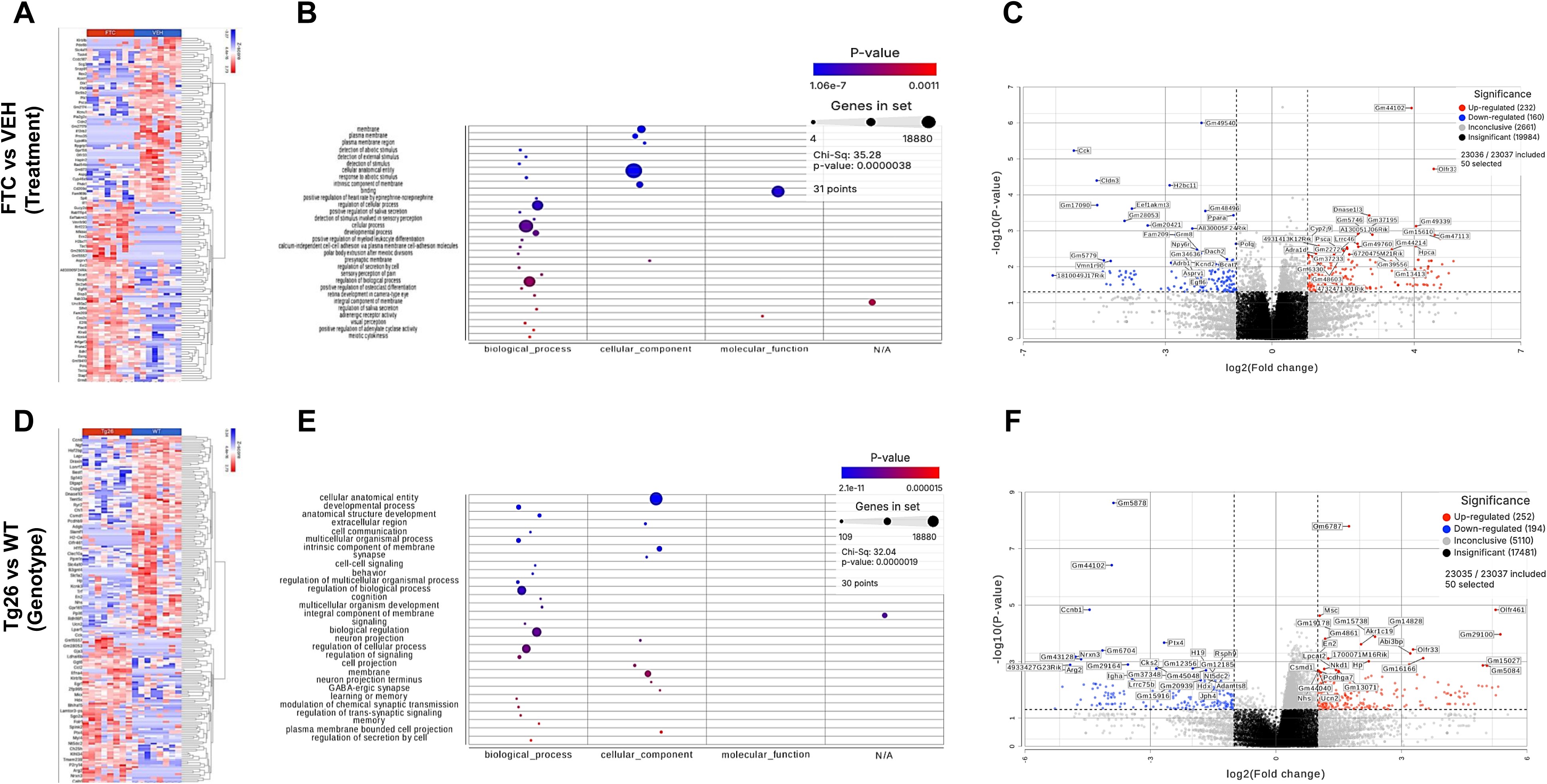
Impact of drug treatments (VEH vs FTC for both genotypes combined) or HIV background (WT vs. Tg26 for both drug treatment groups combined) on differentially regulated genes in skeletal muscle: **(A)** Heatmap shows genes (columns) and comparison groups (rows; each row represents one mouse), where color intensity reflects the Z-score, with blue indicating downregulation and red indicating upregulation **(B)** Gene Ontology plots of protein coding genes and biological processes **(C)** Volcano plot: Top 50 highlighted differentially expressed (upregulated and downregulated) genes among the comparison groups based on log2 fold changes. (**D**) Heatmap shows genes (columns) and comparison groups (rows; each row represents one mouse), where color intensity reflects the Z-score, with blue indicating downregulation and red indicating upregulation. (E) Gene Ontology plots of protein coding genes and biological processes (F) Volcano plot: Top 50 differentially expressed (upregulated and downregulated) genes among the comparison groups based on log2 fold changes.

**Supplementary Figure 5:**
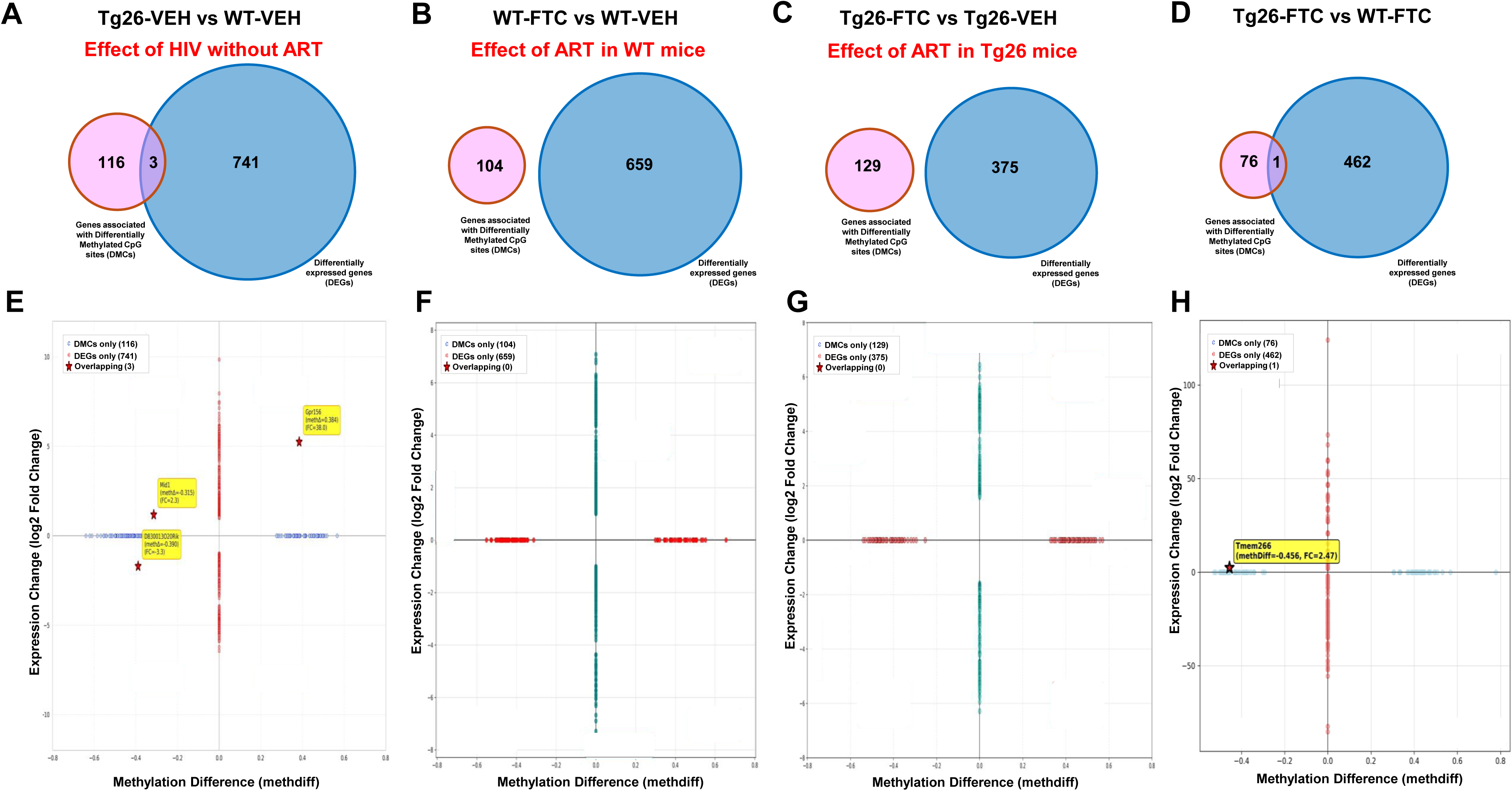
Conjoint analysis of differential methylation and gene expression across four experimental comparison groups. Venn diagrams show the overlap between genes associated with sites of significant differentially methylated CPG sites (DMC) and genes with significant differential expression patterns (DEG) for each comparison group: (A) Tg26-VEH vs WT-VEH, (B) WT-FTC vs WT-VEH, (C) Tg26-FTC vs Tg26-VEH, and (D) Tg26-FTC vs WT-FTC. Numbers inside the Venn diagram circles indicate the count of genes in each category: genes with methylation changes and expression changes only, and genes with both methylation and expression changes (overlapping genes). (E-H) Scatter plots display the relationship between differential methylation (x-axis, methylation difference) and differential gene expression (y-axis, log2 fold change) for the four comparison groups. Each point represents a gene, with colors indicating significance levels and magnitude of changes. Quadrants are labeled to indicate different patterns of coordinated changes: hypermethylation with upregulation (upper right), hypermethylation with downregulation (lower right), hypomethylation with upregulation (upper left), and hypomethylation with downregulation (lower left).

---

**Supplemental Table 1.** g:Profiler analysis: Enriched pathways (p<0.05) associated with DMRs.

| term_id | source | term_name | p value |
| --- | --- | --- | --- |
| **Impact of HIV Phenotype (Tg26-VEH vs WT-VEH)** |  |  |  |
| HP:0011800 | HP | Midface retrusion | 0.007737707 |
| **Impact of ART (WT-FTC vs WT-VEH)** |  |  |  |
| GO:0005834 | GO:CC | heterotrimeric G-protein complex | 0.006568135 |
| GO:1905360 | GO:CC | GTPase complex | 0.008605011 |
| GO:0031234 | GO:CC | extrinsic component of cytoplasmic side of plasma membrane | 0.02808667 |
| GO:0010854 | GO:MF | adenylate cyclase regulator activity | 0.000572384 |
| GO:0010851 | GO:MF | cyclase regulator activity | 0.001667589 |
| GO:0031683 | GO:MF | G-protein beta/gamma-subunit complex binding | 0.004376036 |
| KEGG:04730 | KEGG | Long-term depression | 0.000891911 |
| WP:WP232 | WP | G protein signaling pathways | 0.021490927 |
| **Impact of ART in HIV background (Tg26-FTC vs Tg26-VEH)** |  |  |  |
| CORUM:3003 | CORUM | Smad3 homotrimer complex | 0.024978768 |
| CORUM:1829 | CORUM | SMAD3-VDR complex | 0.049957536 |
| CORUM:2831 | CORUM | TIF1gamma-Smad2-Smad3 complex | 0.049957536 |
| CORUM:3000 | CORUM | Smad2-Smad3 complex | 0.049957536 |
| CORUM:3002 | CORUM | Smad3-Smad4 complex | 0.049957536 |
| CORUM:3713 | CORUM | Smad3-Smad4 complex | 0.049957536 |
| GO:0032906 | GO:BP | transforming growth factor beta2 production | 0.007779233 |
| GO:0032909 | GO:BP | regulation of transforming growth factor beta2 production | 0.007779233 |
| GO:0010586 | GO:BP | miRNA metabolic process | 0.013857179 |
| GO:0071636 | GO:BP | positive regulation of transforming growth factor beta production | 0.04975642 |
| KEGG:04310 | KEGG | Wnt signaling pathway | 0.010132046 |
| WP:WP1763 | WP | Mechanisms associated with pluripotency | 0.01895383 |
| **ART-treated Tg26 vs. WT mice** |  |  |  |
| TF:M07461_1 | TF | Factor: KLF; motif: GGGNGGGG; match class: 1 | 0.034922593 |

**Supplemental Table 2.**
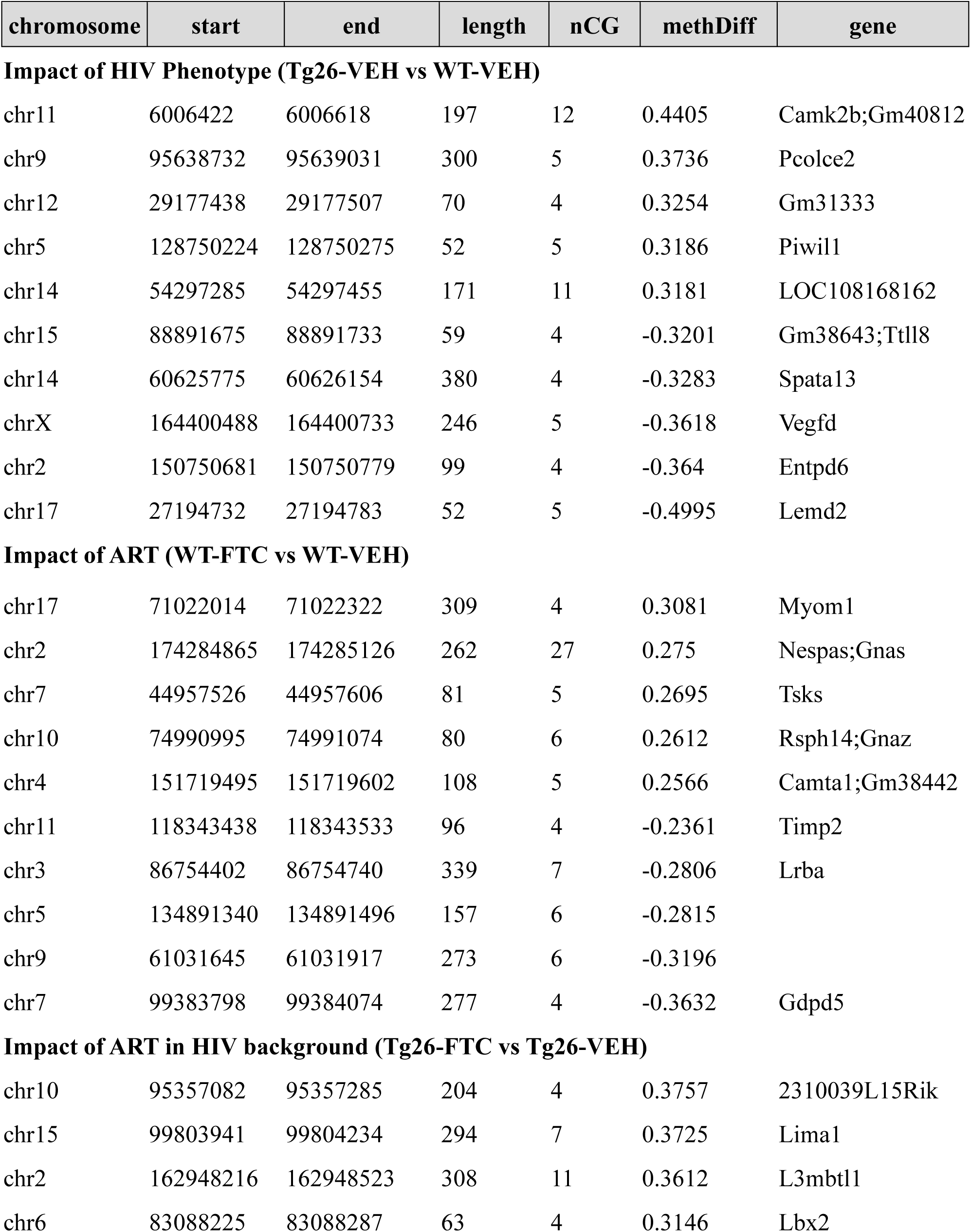
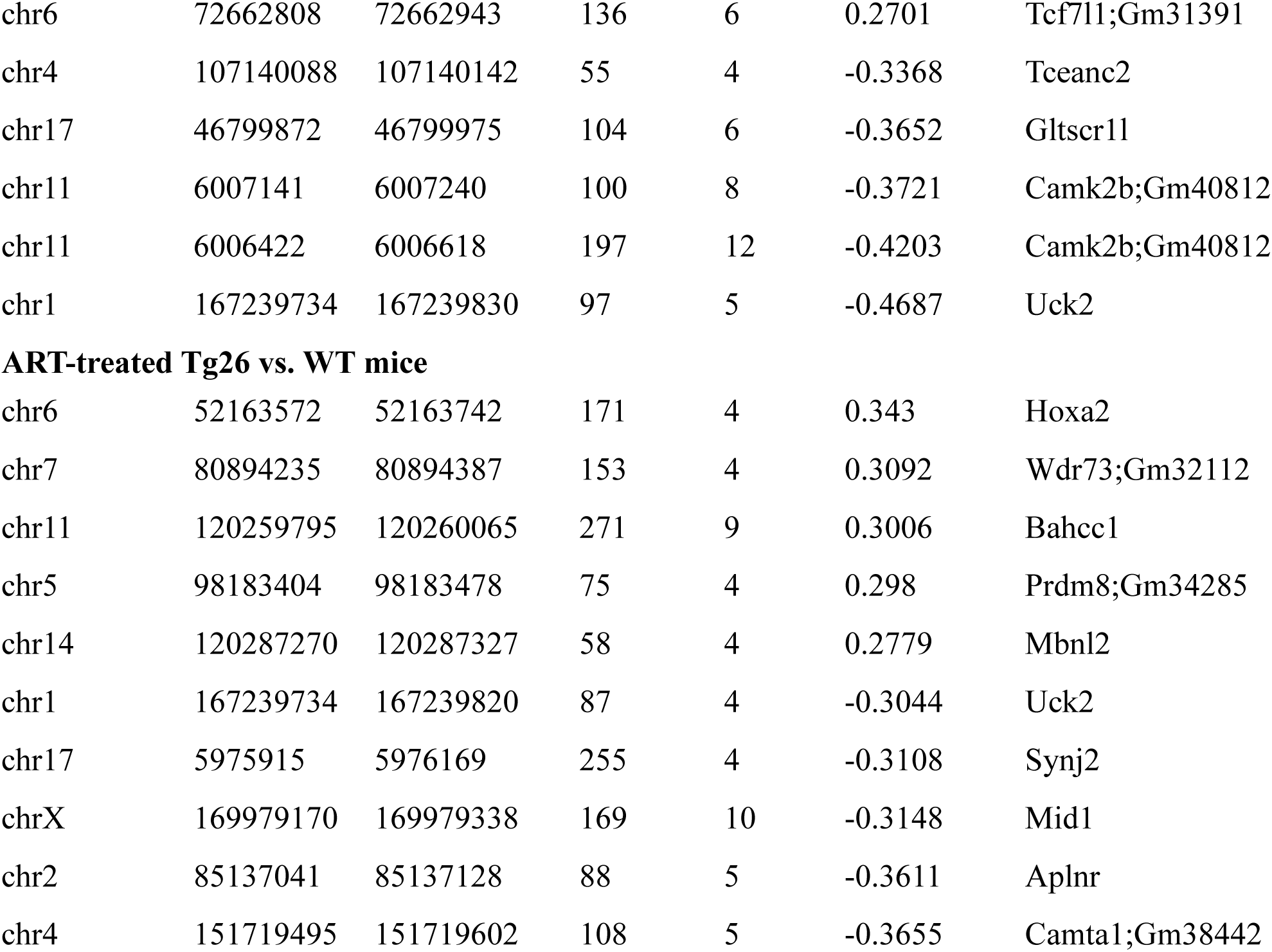
Top 5 differentially (hyper/hypo) methylated genes associated with DMRs.

**Supplemental Table 3.**
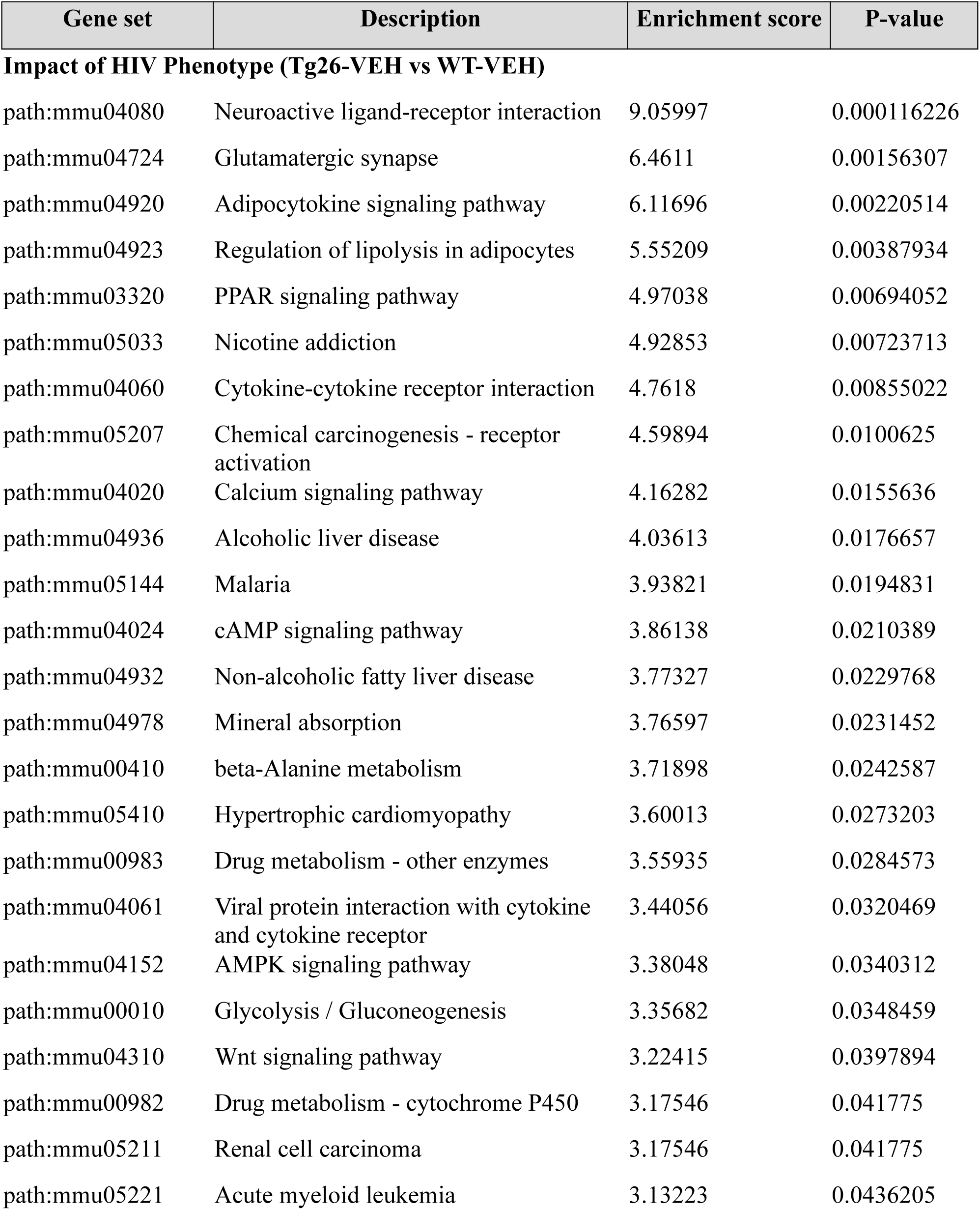
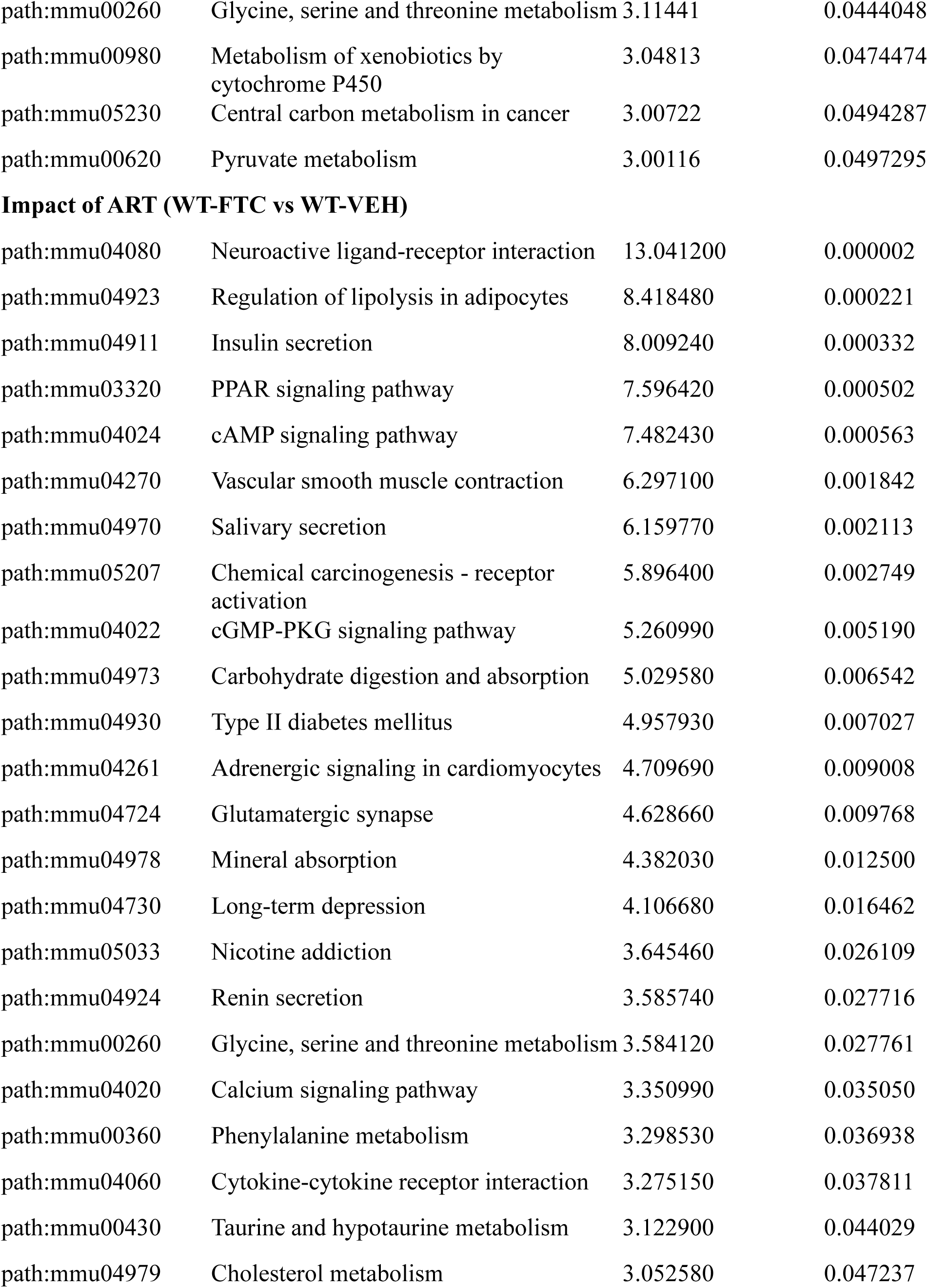
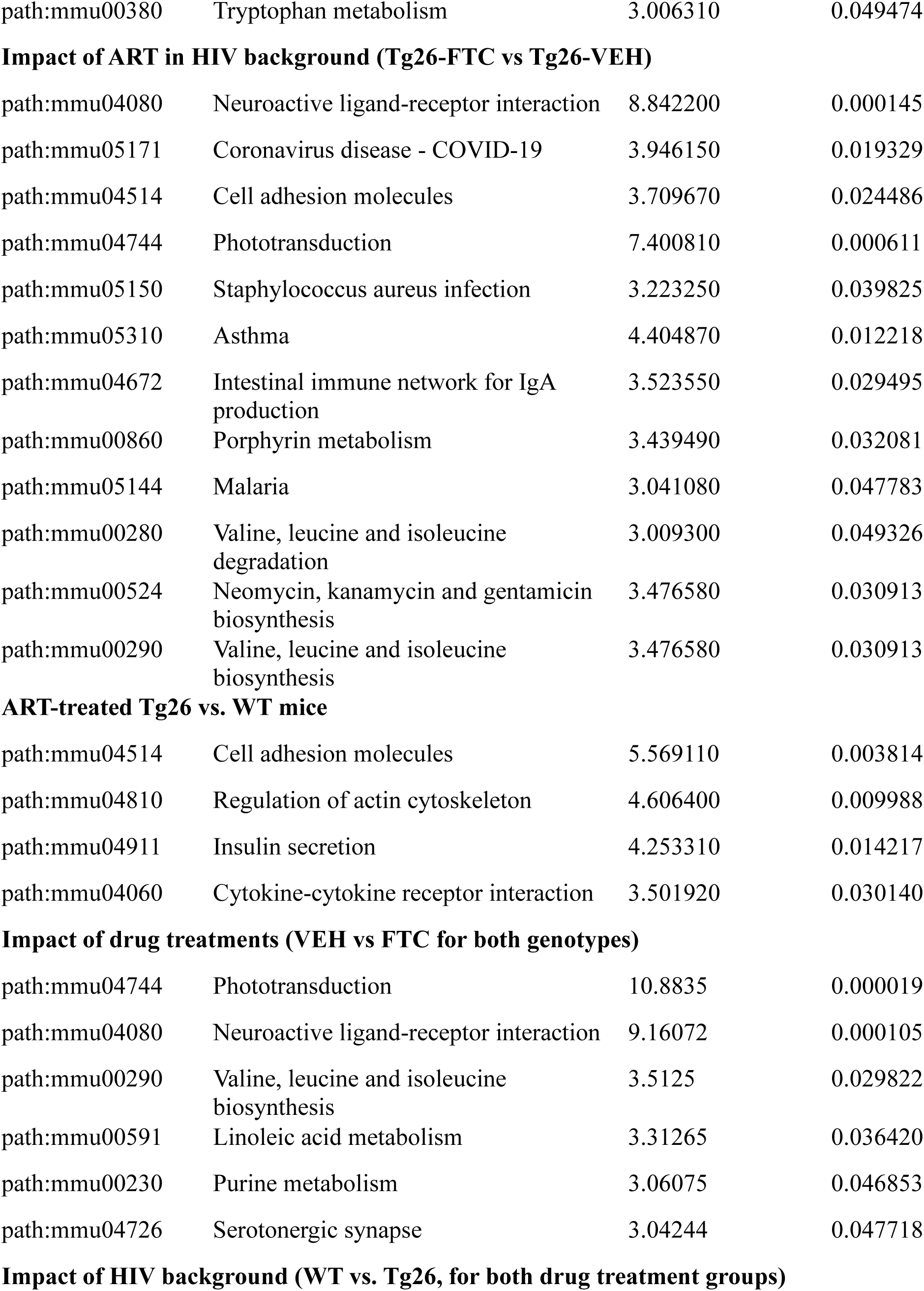
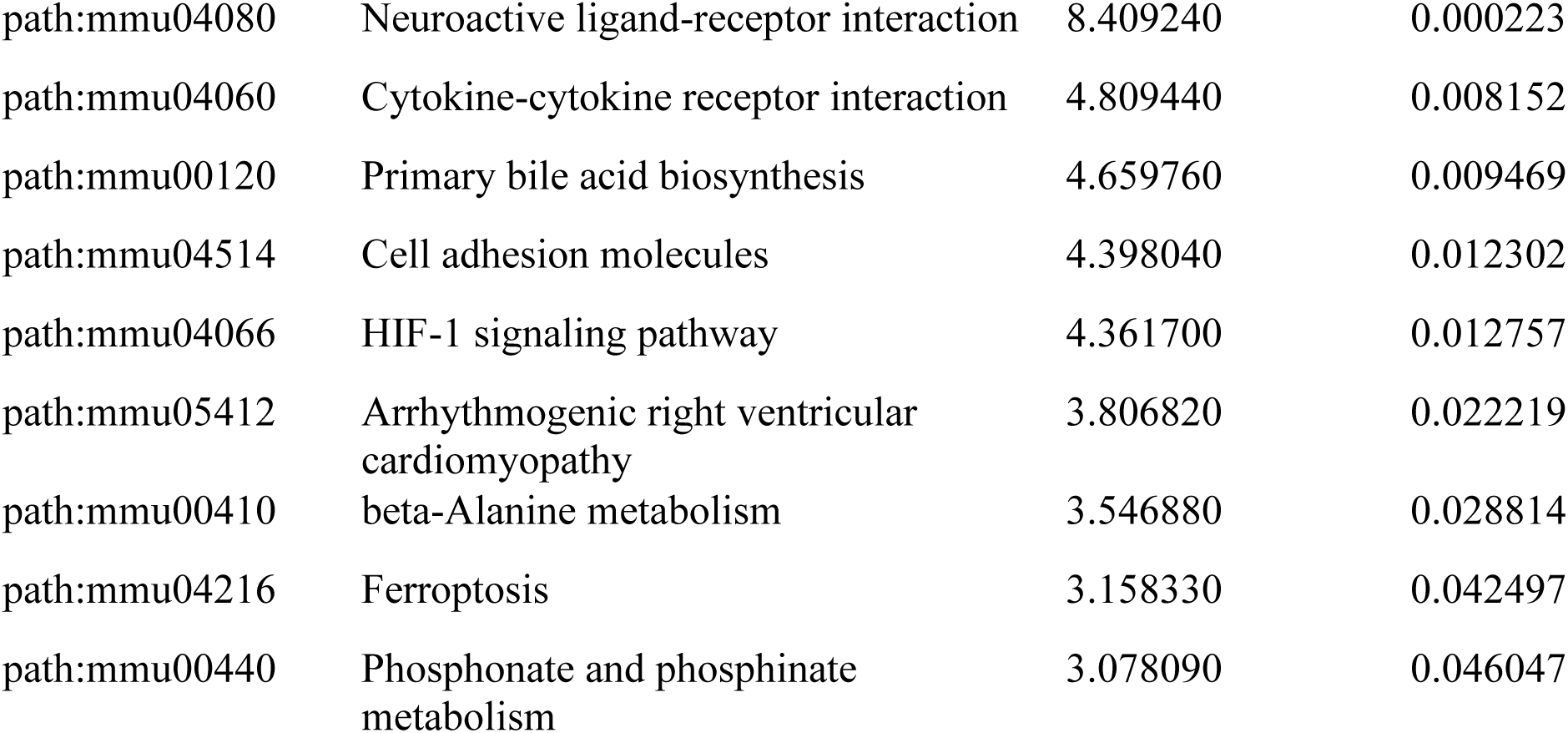
Gene Set Enrichment Analysis (GSEA) of differentially expressed genes (p<0.05, enrichment score ≥3.0)

| Gene set | Description | Enrichment score | P-value |
| --- | --- | --- | --- |
| **Impact of HIV Phenotype (Tg26-VEH vs WT-VEH)** |  |  |  |
| path:mmu04080 | Neuroactive ligand-receptor interaction | 9.05997 | 0.000116226 |
| path:mmu04724 | Glutamatergic synapse | 6.4611 | 0.00156307 |
| path:mmu04920 | Adipocytokine signaling pathway | 6.11696 | 0.00220514 |
| path:mmu04923 | Regulation of lipolysis in adipocytes | 5.55209 | 0.00387934 |
| path:mmu03320 | PPAR signaling pathway | 4.97038 | 0.00694052 |
| path:mmu05033 | Nicotine addiction | 4.92853 | 0.00723713 |
| path:mmu04060 | Cytokine-cytokine receptor interaction | 4.7618 | 0.00855022 |
| path:mmu05207 | Chemical carcinogenesis - receptor activation | 4.59894 | 0.0100625 |
| path:mmu04020 | Calcium signaling pathway | 4.16282 | 0.0155636 |
| path:mmu04936 | Alcoholic liver disease | 4.03613 | 0.0176657 |
| path:mmu05144 | Malaria | 3.93821 | 0.0194831 |
| path:mmu04024 | cAMP signaling pathway | 3.86138 | 0.0210389 |
| path:mmu04932 | Non-alcoholic fatty liver disease | 3.77327 | 0.0229768 |
| path:mmu04978 | Mineral absorption | 3.76597 | 0.0231452 |
| path:mmu00410 | beta-Alanine metabolism | 3.71898 | 0.0242587 |
| path:mmu05410 | Hypertrophic cardiomyopathy | 3.60013 | 0.0273203 |
| path:mmu00983 | Drug metabolism - other enzymes | 3.55935 | 0.0284573 |
| path:mmu04061 | Viral protein interaction with cytokine and cytokine receptor | 3.44056 | 0.0320469 |
| path:mmu04152 | AMPK signaling pathway | 3.38048 | 0.0340312 |
| path:mmu00010 | Glycolysis / Gluconeogenesis | 3.35682 | 0.0348459 |
| path:mmu04310 | Wnt signaling pathway | 3.22415 | 0.0397894 |
| path:mmu00982 | Drug metabolism - cytochrome P450 | 3.17546 | 0.041775 |
| path:mmu05211 | Renal cell carcinoma | 3.17546 | 0.041775 |
| path:mmu05221 | Acute myeloid leukemia | 3.13223 | 0.0436205 |
| path:mmu00260 | Glycine, serine and threonine metabolism | 3.11441 | 0.0444048 |
| path:mmu00980 | Metabolism of xenobiotics by cytochrome P450 | 3.04813 | 0.0474474 |
| path:mmu05230 | Central carbon metabolism in cancer | 3.00722 | 0.0494287 |
| path:mmu00620 | Pyruvate metabolism | 3.00116 | 0.0497295 |
| **Impact of ART (WT-FTC vs WT-VEH)** |  |  |  |
| path:mmu04080 | Neuroactive ligand-receptor interaction | 13.041200 | 0.000002 |
| path:mmu04923 | Regulation of lipolysis in adipocytes | 8.418480 | 0.000221 |
| path:mmu04911 | Insulin secretion | 8.009240 | 0.000332 |
| path:mmu03320 | PPAR signaling pathway | 7.596420 | 0.000502 |
| path:mmu04024 | cAMP signaling pathway | 7.482430 | 0.000563 |
| path:mmu04270 | Vascular smooth muscle contraction | 6.297100 | 0.001842 |
| path:mmu04970 | Salivary secretion | 6.159770 | 0.002113 |
| path:mmu05207 | Chemical carcinogenesis - receptor activation | 5.896400 | 0.002749 |
| path:mmu04022 | cGMP-PKG signaling pathway | 5.260990 | 0.005190 |
| path:mmu04973 | Carbohydrate digestion and absorption | 5.029580 | 0.006542 |
| path:mmu04930 | Type II diabetes mellitus | 4.957930 | 0.007027 |
| path:mmu04261 | Adrenergic signaling in cardiomyocytes | 4.709690 | 0.009008 |
| path:mmu04724 | Glutamatergic synapse | 4.628660 | 0.009768 |
| path:mmu04978 | Mineral absorption | 4.382030 | 0.012500 |
| path:mmu04730 | Long-term depression | 4.106680 | 0.016462 |
| path:mmu05033 | Nicotine addiction | 3.645460 | 0.026109 |
| path:mmu04924 | Renin secretion | 3.585740 | 0.027716 |
| path:mmu00260 | Glycine, serine and threonine metabolism | 3.584120 | 0.027761 |
| path:mmu04020 | Calcium signaling pathway | 3.350990 | 0.035050 |
| path:mmu00360 | Phenylalanine metabolism | 3.298530 | 0.036938 |
| path:mmu04060 | Cytokine-cytokine receptor interaction | 3.275150 | 0.037811 |
| path:mmu00430 | Taurine and hypotaurine metabolism | 3.122900 | 0.044029 |
| path:mmu04979 | Cholesterol metabolism | 3.052580 | 0.047237 |
| path:mmu00380 | Tryptophan metabolism | 3.006310 | 0.049474 |
| **Impact of ART in HIV background (Tg26-FTC vs Tg26-VEH)** |  |  |  |
| path:mmu04080 | Neuroactive ligand-receptor interaction | 8.842200 | 0.000145 |
| path:mmu05171 | Coronavirus disease - COVID-19 | 3.946150 | 0.019329 |
| path:mmu04514 | Cell adhesion molecules | 3.709670 | 0.024486 |
| path:mmu04744 | Phototransduction | 7.400810 | 0.000611 |
| path:mmu05150 | Staphylococcus aureus infection | 3.223250 | 0.039825 |
| path:mmu05310 | Asthma | 4.404870 | 0.012218 |
| path:mmu04672 | Intestinal immune network for IgA production | 3.523550 | 0.029495 |
| path:mmu00860 | Porphyrin metabolism | 3.439490 | 0.032081 |
| path:mmu05144 | Malaria | 3.041080 | 0.047783 |
| path:mmu00280 | Valine, leucine and isoleucine degradation | 3.009300 | 0.049326 |
| path:mmu00524 | Neomycin, kanamycin and gentamicin biosynthesis | 3.476580 | 0.030913 |
| path:mmu00290 | Valine, leucine and isoleucine biosynthesis | 3.476580 | 0.030913 |
| **ART-treated Tg26 vs. WT mice** |  |  |  |
| path:mmu04514 | Cell adhesion molecules | 5.569110 | 0.003814 |
| path:mmu04810 | Regulation of actin cytoskeleton | 4.606400 | 0.009988 |
| path:mmu04911 | Insulin secretion | 4.253310 | 0.014217 |
| path:mmu04060 | Cytokine-cytokine receptor interaction | 3.501920 | 0.030140 |
| **Impact of drug treatments (VEH vs FTC for both genotypes)** |  |  |  |
| path:mmu04744 | Phototransduction | 10.8835 | 0.000019 |
| path:mmu04080 | Neuroactive ligand-receptor interaction | 9.16072 | 0.000105 |
| path:mmu00290 | Valine, leucine and isoleucine biosynthesis | 3.5125 | 0.029822 |
| path:mmu00591 | Linoleic acid metabolism | 3.31265 | 0.036420 |
| path:mmu00230 | Purine metabolism | 3.06075 | 0.046853 |
| path:mmu04726 | Serotonergic synapse | 3.04244 | 0.047718 |
| **Impact of HIV background (WT vs. Tg26, for both drug treatment groups)** |  |  |  |
| path:mmu04080 | Neuroactive ligand-receptor interaction | 8.409240 | 0.000223 |
| path:mmu04060 | Cytokine-cytokine receptor interaction | 4.809440 | 0.008152 |
| path:mmu00120 | Primary bile acid biosynthesis | 4.659760 | 0.009469 |
| path:mmu04514 | Cell adhesion molecules | 4.398040 | 0.012302 |
| path:mmu04066 | HIF-1 signaling pathway | 4.361700 | 0.012757 |
| path:mmu05412 | Arrhythmogenic right ventricular<br>cardiomyopathy | 3.806820 | 0.022219 |
| path:mmu00410 | beta-Alanine metabolism | 3.546880 | 0.028814 |
| path:mmu04216 | Ferroptosis | 3.158330 | 0.042497 |
| path:mmu00440 | Phosphonate and phosphinate<br>metabolism | 3.078090 | 0.046047 |

